# Structural insights into target detection by the *S. marcescens* type III CRISPR complex and its deployment in SNP identification

**DOI:** 10.64898/2026.03.30.715313

**Authors:** Calvin C. Perdigao, Luqman O. Ajisafe, Anju T. Sunny, Si Wu, Terje Dokland, Jack A. Dunkle

## Abstract

Type III CRISPR systems utilize a complex containing Cas10, additional Cas proteins and a crRNA to detect foreign transcripts. Upon detection, Cas10 synthesizes cyclic oligoadenylates (cOA), signaling molecules that coordinate interference by stimulating downstream enzymes with DNase, RNase, protease or other activities. Type III systems are among the most abundant CRISPR systems in prokaryotes and understanding the structure-function relationships that control transcript detection and cOA synthesis will advance the understanding of the broader physiological roles of these systems. Type III systems possess properties well-suited to their deployment as molecule diagnostics: specific detection and activation of a cascade of multi-turnover enzymatic reactions that can be harnessed for signal generation. We determined that *Serratia marcescens* Cas10-Csm (SmCas10-Csm) synthesizes predominantly cA_3_ molecules and this synthesis is sensitive to mismatches in the crRNA-target RNA duplex adjacent to Cas10. We determined the structure of SmCas10-Csm unbound and bound to target RNA identifying conformational changes associated with target binding. We demonstrate that SmCas10-Csm can distinguish between single nucleotide polymorphisms that occur in the human *HBB* transcript that are associated with sickle cell disease indicating an additional role for type III CRISPR systems in point-of-care diagnostics in low-resource settings.

## Introduction

CRISPR systems, which are present in a large fraction of prokaryotic genomes, use a crRNA bound to a Cas (<u>C</u>RISPR-<u>as</u>sociated) protein, or a multi-protein complex of Cas proteins, to detect and interfere with foreign genetic elements (1–4). CRISPR systems are grouped into two classes, seven types and many subtypes (5). Class 2 systems, such as Cas9 and Cas12 use a single Cas protein to carry out interference while class 1 systems use a multi-protein complex for interference (5).

Type III CRISPR systems are multi-protein complexes that contain the Cas10 effector protein capable of generating a second messenger molecule polymerized from ATP, termed cyclic oligoadenylate (cOA). Use of a second messenger molecule to coordinate interference is a feature unique to type III systems and gives rise to a modularity: the crRNA-containing Cas10 complex senses the presence of a foreign transcript but interference, in part, is carried out by enzymes activated by cOA allowing evolution, gene acquisition and fusion to generate diverse biochemical activities responding to cOA. Recent results highlight this phenomenon with the identification of RNase, DNase and protease activities stimulated by cOA (6–11). Additionally, membrane depolarization controlled by cOA sensing has been reported (12–14).

Many bacterial genomes contain multiple CRISPR systems including that of *Serratia marcescens* ATCC 39006 (recently renamed *Prodigiosinella aquatilis*) which possesses types I-E, I-F and III-A CRISPR systems (Fig. 1) (11,15). Type I-E and I-F CRISPR systems both sense foreign DNA by its ability to base-pair to crRNA in complex with Cas proteins while the III-A system senses foreign RNA via base-pairing to the crRNA-Cas complex (3). A justification for the presence of systems which sense both DNA and RNA within the same genome is emerging. Jumbo phage, defined as phage with a genome of > 200 kb, can encapsulate their genome within a proteinaceous structure termed a phage nucleus which protects it from interference by CRISPR systems that sense and cleave DNA (16,17). The transcripts of these jumbo phage, however, are still sensitive to detection by type III-A systems (17). Recently it was shown that the type III-A CRISPR system of *S. marcescens* senses jumbo phage transcripts activating cOA synthesis by Cas10. These cOA bind to and stimulate the DNase activity of NucC, leading to degradation of the host genome to block phage replication in the wider population via an abortive infection mode (11).

**Figure 1.**
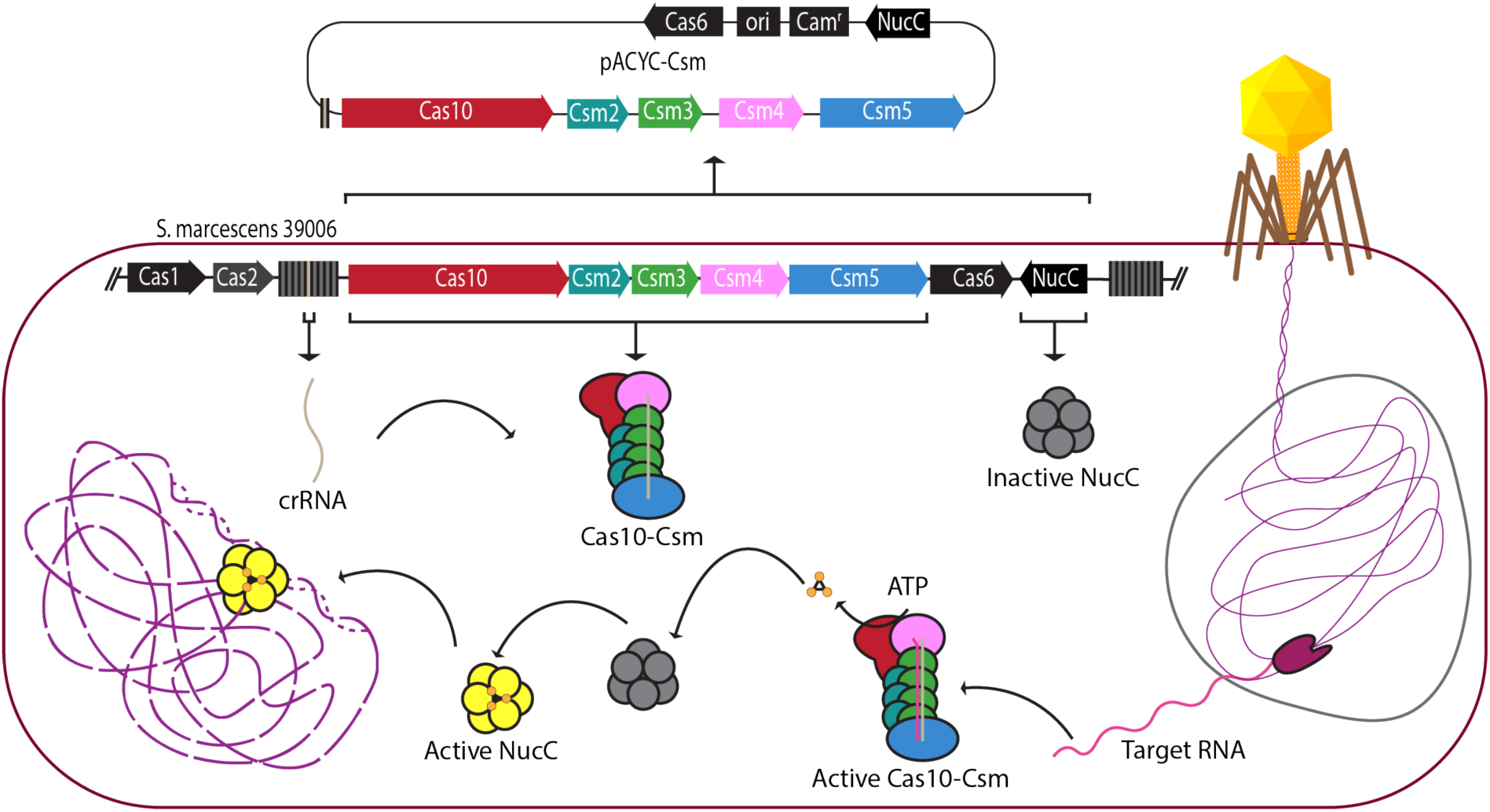
Overview of interference mediated by the type III CRISPR system of *S. marcescens*. The *S. marcescens* type III CRISPR system detects bacteriophage transcripts to activate synthesis of cyclic oligoadenylates (cOA). Jumbo phage transcript detection occurs despite the bacteriophage genome’s encapsulation in a phage nucleus (grey line). COA bind to the NucC endonuclease stimulating its DNase activity. Activated NucC degrades the bacterial genome (purple dashed line) blocking bacteriophage spread by an abortive infection mechanism.

While the evolutionary justification for a type III-A system in *S. marcescens* has now been established, the precise architecture of this system, such as the oligomeric state of its Cas10-Csm complex, is not known, and neither have the conformational changes that occur upon target RNA binding been described. In this study, we determined the cryo-EM structures of the *S. marcescens* Cas10-Csm unbound and bound to target RNA to address these issues. We also determined that *S. marcescens* Cas10-Csm synthesizes 3-mer and 4-mer lengths of cOA and identified the effect of mismatches between the crRNA and target RNA on cOA synthesis. This structure-function information will facilitate the use of *S. marcescens* as a model system for studying the dynamic events and mechanisms of interference against jumbo phages.

The development of molecular diagnostics based upon detection of a nucleic acid by a CRISPR system has recently garnered much interest. While there are many other powerful methodologies for detecting the presence of a nucleic acid, techniques such as microarrays, DNA sequencing or PCR require skilled technicians and expensive laboratory equipment that are challenging to deploy in low resource settings. Tools based on Cas12 or Cas13 systems have shown promise as point-of-care tests or field tests for detecting viruses, pathogenic bacteria and genotyping human DNA (18–21). Cas10 containing CRISPR systems have also been employed as molecular diagnostics (8,22–25). Cas10 systems have an advantage over Cas12 and Cas13 systems in that target RNA binding activates cOA synthesis by Cas10. Many cOA-activated enzymes with distinct activities have been described (6–10,12–14,26); therefore, RNA detection and subsequent cOA synthesis can be linked to many different enzymatic activities for read-out. To our knowledge, detection of a human SNP by Cas10-Csm has not previously been demonstrated. We show that *S. marcescens* Cas10-Csm can be re-purposed to detect the most common SNP associated with the formation of sickle cell disease.

## Materials and Methods

### Generation of pACYC-Csm and its site-directed mutants

The plasmid pACYC-Csm was generated by Genscript using gene synthesis. Briefly, the sequences present in the type III-A CRISPR locus of *Serratia marcescens ATCC 39006* were inserted into a pACYC backbone. This includes *spc5* flanked by an associated repeat sequence to form a mini-repeat spacer array and the genes for *cas6*, *cas10*, *csm2*, *csm3*, *csm4*, *csm5* and *nucC*. The plasmid map is given in Figure S1. Catalytically dead site-directed mutants were generated for the Cas10 Palm2 domain (dPalm2, G616-D619 to A616-A619), NucC (E114A or D83Q) and Csm3 (dCsm3, D34A) using New England Biolabs Q5 mutagenesis or by inserting gene strands to the plasmid as indicated in Table S1. Cas10-Csm complexes encoding crRNA targeting the *HBB* transcript of human β-globin were generated as follows: gene synthesis was used to create the repeat-spacer array and it was inserted into pACYC-Csm-dCsm3 using NcoI and PspXI restriction endonuclease sites.

### In vivo interference assays

Cells harboring pACYC-Csm were electroporated with pTarget and outgrown for 1 hour at 37 °C in SOC media. Serial dilutions were plated on LB agar containing 17 μg/mL chloramphenicol (Cam) and 50 μg/mL ampicillin (Amp). An incubation was performed at 37 °C for 11 hours and then colonies were counted.

### Purification of Cas10-Csm

An overnight culture of *E. coli* BL21 AI harboring pACYC-Csm or a site-directed mutant thereof was diluted 100-fold into 1 L of LB media containing 34 µg/mL Cam. The culture was incubated at 37°C with 180 rpm shaking until the OD_600_ reached 0.6-0.8. After 30 minutes on ice, recombinant expression was induced with 0.02%-0.2% (w/v) L-arabinose and continued overnight at 18°C. Cells were harvested by centrifugation and pellets stored at - 20°C until use. For Cas10-Csm complexes carrying a crRNA derived from *spc5* (which targets the *DotA* transcript), cells were resuspended in phosphate lysis buffer (Table S2). The cell suspension was incubated on ice for 1 hour and then lysed by sonication. Lysate was cleared by centrifugation and filtered with a 0.8 µm filter. Immobilized metal affinity chromatography (IMAC) was carried out using phosphate wash buffer and phosphate elution buffer (Table S2). For Cas10-Csm complexes carrying a crRNA targeting the *HBB* transcript, cells were resuspended in Tris lysis buffer (Table S2). Lysis and clarification of the lysate was performed as described above and IMAC was carried out using Tris wash buffer and Tris elution buffer (Table S2). For all Cas10-Csm complexes, fractions containing the complex were pooled and layered on 5-20% (w/v) sucrose gradient (made with 50 mM Tris HCl pH 8.0, 150 mM NaCl, and 5% (v/v) glycerol). Then ultracentrifugation on Beckman SW-32Ti rotor was performed at 30000 RPM for 41 hours at 4°C. Fractions containing the complex were combined, buffer exchanged to storage buffer (Table S2), concentrated with a 10 KDa MWCO Amicon Ultra centrifugal filter and flash-frozen for storage at −80°C.

### Analysis of crRNA

CrRNA co-purifying with Cas10-Csm was extracted with a 1:1 (v/v) ratio of phenol chloroform-isoamyl alcohol (25:24:1) twice, followed by one extraction with 1 volume chloroform. It was precipitated with isopropyl alcohol and resuspended in 20 μL water. Radiolabeling of crRNA was performed using γ-^32^ATP (3000 Ci/mmol) and T4 Polynucleotide kinase at 37 °C for 1 hour. Next, crRNA was mixed 1:1 with formamide dye and heated at 70°C for 2 minutes before loading onto a 12% acrylamide, 8 M urea gel. An Invitrogen Decade Marker was used to generate labeled RNA markers to estimate the crRNA size. Electrophoresis was carried out at 50 W for ∼90 min. Phosphor imaging was performed using a Typhoon FLA 7000 from GE Healthcare.

### LC-MS/MS Analysis

Digests of the Cas10-Csm ultracentrifugation fraction were diluted to a final concentration of 125 ng/µL in 10 µL of HPLC-grade water. A 1 µL aliquot of the resulting solution was then analyzed by LC-MS/MS using a Vanquish Neo UPLC system coupled with an Orbitrap Ascend mass spectrometer (Thermo Fisher Scientific). The sample was injected onto a PepMap™ Neo Trap Cartridge (5 µm beads, 300 µm x 5 mm, Thermo Fisher Scientific) before being separated on a PepMap™ Neo C18 analytical column (2 µm beads, 75 µm x 150 mm, Thermo Fisher Scientific). Mobile phase A (MPA) consisted of 0.1% formic acid in water, while mobile phase B (MPB) consisted of 0.1% formic acid in 80% acetonitrile. A 40-minute gradient from 2% to 40% MPB was applied. Data-dependent acquisition (DDA) was used for LC-MS/MS analysis. The full MS scan was acquired over a m/z range of 375-1500, with a resolution of 120,000 at m/z = 200. The AGC target for MS scans was set to 4 × 10^5^, and the maximum ion injection time is 251 ms. Precursor ions with charge states between 2 and 7 were selected for higher-energy collisional dissociation (HCD) fragmentation with a cycle time of 3 s. The resolution of MS/MS scans was 60,000 at m/z = 200. The AGC target for MS/MS scans was set to 5 × 10^4^, and the maximum ion injection time was set to 123 ms. The normalized collision energy for HCD was set to 25%, with an isolation width of 2 m/z. Dynamic exclusion was set to 60 seconds. Mass spectrometry data were processed using Proteome Discoverer 3.1 (Thermo Fisher Scientific). Database search was performed using a customized protein database that combined the *E. coli* proteome (4217 sequences retrieved from UniParc, released 2024_09) with the sequences of 5 proteins of interest from the SmCas10-Csm complex. Proteins were identified at a 1% False Discovery Rate (FDR) with two uniquely identified peptides.

### Detection of cOA by mass spectrometry

Biosynthesis of cOA was performed in reaction buffer 1 (Table S2) with 500 μM ATP, 200 nM target RNA (*DotA*), 200 nM Cas10-Csm. Reactions proceeded overnight at 37°C. COAs were extracted with phenol chloroform-isoamyl alcohol (25:24:1) as described above. Samples were then zip tipped with C18 tips and subjected to MALDI-MS using a matrix of 50 mg/mL 3-HPA in 50 % acetonitrile and 50 mg/mL tri-ammonium citrate.

### COA synthesis measured by the malachite green assay

Reactions contained 250 μM ATP, 200 nM target RNA (*DotA)*, 100 nM Cas10-Csm and 0.05 units of TIPP (New England Biolabs) in reaction buffer 2. A one-hour incubation at 37 °C was performed then cOA synthesis was quenched by a 75 °C incubation for 10 minutes. The TIPP enzyme converts pyrophosphate products of cOA synthesis to phosphate. The Malachite Green Phosphate Detection Kit (Thermo Fisher Scientific) and a Bio-Tek Synergy H1 plate reader set to read at 620 nm were used to quantify phosphate. This assay was performed with cognate *DotA* target RNA and a panel of *DotA* target RNAs with single mismatches to crRNA. The panel was generated by in vitro transcription from DNA duplexes described in Table S1 using the Promega RiboMAX kit following the manufacturer’s instructions.

### Detection of ^32^P-labeled cOA by TLC

Reactions generating ^32^P-labeled cOA contained 500 μM unlabeled ATP, 0.1 μL α-^32^P-ATP 3000 Ci/mmol (Revvity), 100 nM Cas10-Csm and 100 nM target RNA (*DotA*) in reaction buffer 2. Incubation at 37°C for one hour was performed followed by a 95°C two minute quench. Separation of the ^32^P-cOA product from the α-^32^P-ATP reactant was performed by TLC as described in (27).

### Structure determination of SmCas10-Csm by cryo-EM

Cas10-Csm was used at a concentration of 3.75 µM in EM buffer (Table S2) and combined with ssRNA-d02 target RNA at the same concentration for the protein-RNA complex. The samples were applied to glow-discharged copper grids with holey carbon supports (Quantifoil R2/1) and plunge frozen in liquid ethane using a Vitrobot Mark IV. Cryo-EM movies were collected using EPU on a Thermo Fisher Glacios 2 electron microscope with a Falcon 4i direct detector. The movies were processed and single-particle reconstruction was performed in CryoSPARC 4.4.0. For the Cas10-Csm-RNA complex, 10,388 movies were subjected to patch motion correction and patch CTF fitting. The 534 best exposures were used for blob particle picking, resulting in 36,058 particles that were subjected to 2D classification to generate templates for template-based particle picking. Iterative 2D classification with rejection of junk particles was performed, yielding 129,762 particles that were used for ab initio reconstruction. Homogeneous and non-uniform refinement were performed resulting in a reconstruction at 3.84 Å resolution (FSC=0.143, Table S4 Fig S4). A coordinate model was constructed by docking AlphaFold 3-generated coordinates into the map in ChimeraX followed by rigid-body refinement in Phenix. Each protein chain was used as a rigid group except for Cas10 which consisted of four rigid bodies, one for each domain of the protein.

Reconstruction of target-free Cas10-Csm was performed using 10,231 movies similarly processed in CryoSPARC. Single-particle reconstruction was performed in CryoSPARC as described above resulting in a reconstruction at 4.47Å resolution (FSC=0.143, Table S4, Fig. S4). AlphaFold 3 generated coordinates were placed in the map using ChimeraX and rigid body fitting was performed in Phenix as described above.

### Multiple sequence alignments

The structure-based multiple sequence alignments given in Figure 7 were generated by superposition of the secondary structure elements of either the Palm2 domain of Cas10, domain 4 of Cas10 or Csm3. Coordinates are given by 9zs4 (*S. marcescens*), 6ifu (*S. thermophilus*), 6mur (*T. onnurineus*), 6xn4 (*L. lactis*), 9g9c (*E. italicus*) and 7v02 (*S. epidermidis*). The Chimera Match->Align tool was used to generate fasta files which were rendered by ESPript (28–30).

### Detection of SNPs by Cas10-Csm

The detection of RNA with the NM_000518.5:c.20A>T (A20T) single nucleotide polymorphism found in the human β-globin allele was performed in the following manner. RNAs mimicking A20T or wild type *HBB* were synthesized by Horizon Discovery (Table S1). A fluorescence assay was established to read-out the presence of the RNA. Purified Cas10-Csm-dCsm3 (250 nM) bound to a crRNA targeting A20T was incubated with 500 μM ATP in reactio buffer 3 (Table S2). To measure the limit-of-detection, A20T RNA or *HBB* RNA varying from 0-1000 nM was added. A 2-minute incubation at 37 °C produced cOA. A 95 °C incubation for 10 minutes was performed to quench cOA synthesis and the reaction was allowed to return to room temperature. Next, 125 nM of a reporter DNA duplex labeled with 5’FAM on one strand and 3’IBFQ quencher on the complementary strand was added to the reaction along with 250 nM NucC. Fluorescence arising from cOA stimulated cleavage of the fluorescent DNA reporter was measured on a BioTek Synergy H1 plate reader using wavelengths: 485 nm excitation and 520 nm emission.

The ability of Cas10-Csm to discriminate between A20T RNA and *HBB* RNA was also tested in contrived samples. These assays were performed as above except 100 ng total human liver RNA (Clontech Laboratories) was added to the reactions with either 5 nM of A20T RNA, 5 nM of *HBB* RNA or 2.5 nM of each.

## Results

### A recombinant interference assay for *S. marcescens* CRISPR*-*Cas10

We sought to establish an easily manipulable interference assay based on recombinant expression of the *S. marcescens* type III-A CRISPR-Cas system in *E. coli*. We used gene synthesis to construct the plasmid pACYC-CSM (Figs. 1, S1 and Table S1). The plasmid contains *spc5* which encodes a crRNA*, cas10*, the *csm2-5* genes which in combination with *cas10* are responsible for directing the formation of the Cas10-Csm ribonucleoprotein, *cas6* which encodes the nuclease required for maturation of the 5’-end of crRNA and *nucC*, a cyclic oligoadenylate activated nuclease that participates in interference. *E. coli* BL21 (DE3) cells were transformed with this plasmid and a transformant was collected to establish an *E. coli* strain capable of interference. We also constructed pTarget-Cognate, which encodes a fragment of *dotA* whose transcript is targeted by Cas10-Csm bound to a crRNA derived from *spc5* (Fig. S2). An additional plasmid, pTarget-Non-cognate serves as a negative control in our interference assay. Binding of SmCas10-Csm to cognate target RNA, an RNA that base-pairs to the body of the crRNA but not to the 5’ tag, stimulates interference, while binding of a non-cognate target RNA—an RNA that base-pairs to the body of the crRNA and with the 5’ tag region—is expected to inhibit cOA synthesis (31).

The base-pairing pattern of cognate targets is expected in true bacteriophage transcripts, while the pairing pattern of non-cognate targets is expected for ‘self’ products resulting from transcription of the CRISPR locus in the reverse direction (Fig. 2).

**Figure 2.**
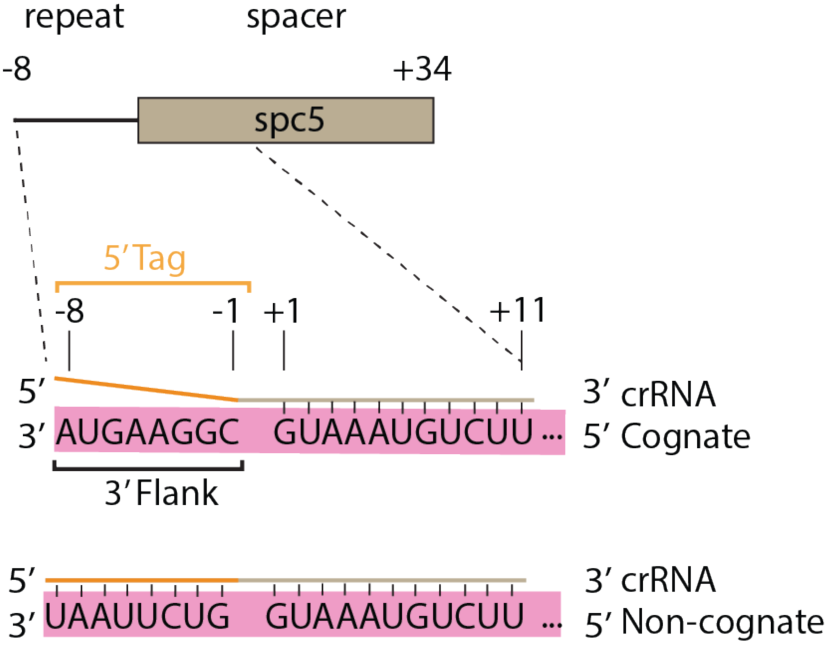
Characteristics of cognate and non-cognate target RNAs. In type III CRISPR systems, interference is stimulated by transcripts which base-pair to the crRNA body, but not to the 5’ tag, and are termed cognate targets. Transcripts which base-pair to the body and the 5’ tag, as would occur from transcription of the anti-sense strand of the CRISPR locus, do not stimulate interference and are termed non-cognate targets.

We carried out our interference assays by introducing pTarget-Cognate to the strain already transformed with pACYC-Csm (Fig. 3A). When these cells are plated on media containing both Cam and Amp antibiotics, the appearance of few or no colonies indicates interference is occurring whereas numerous colonies indicate efficient uptake of the plasmid and a lack of interference (Fig. 3A). As controls, cells carrying pACYC-Csm are plated on Cam containing plates to ensure cell viability and also transformed with pTarget-Non-cognate which should be efficiently taken up (Fig. 2B). When the strain carrying wild type pACYC-Csm was assayed for interference with pTarget-Cognate no transformants were observed but 10^9^ transformants were observed with the pTarget-Non-cognate indicating our assay worked as expected (Fig. 3B).

**Figure 3.**
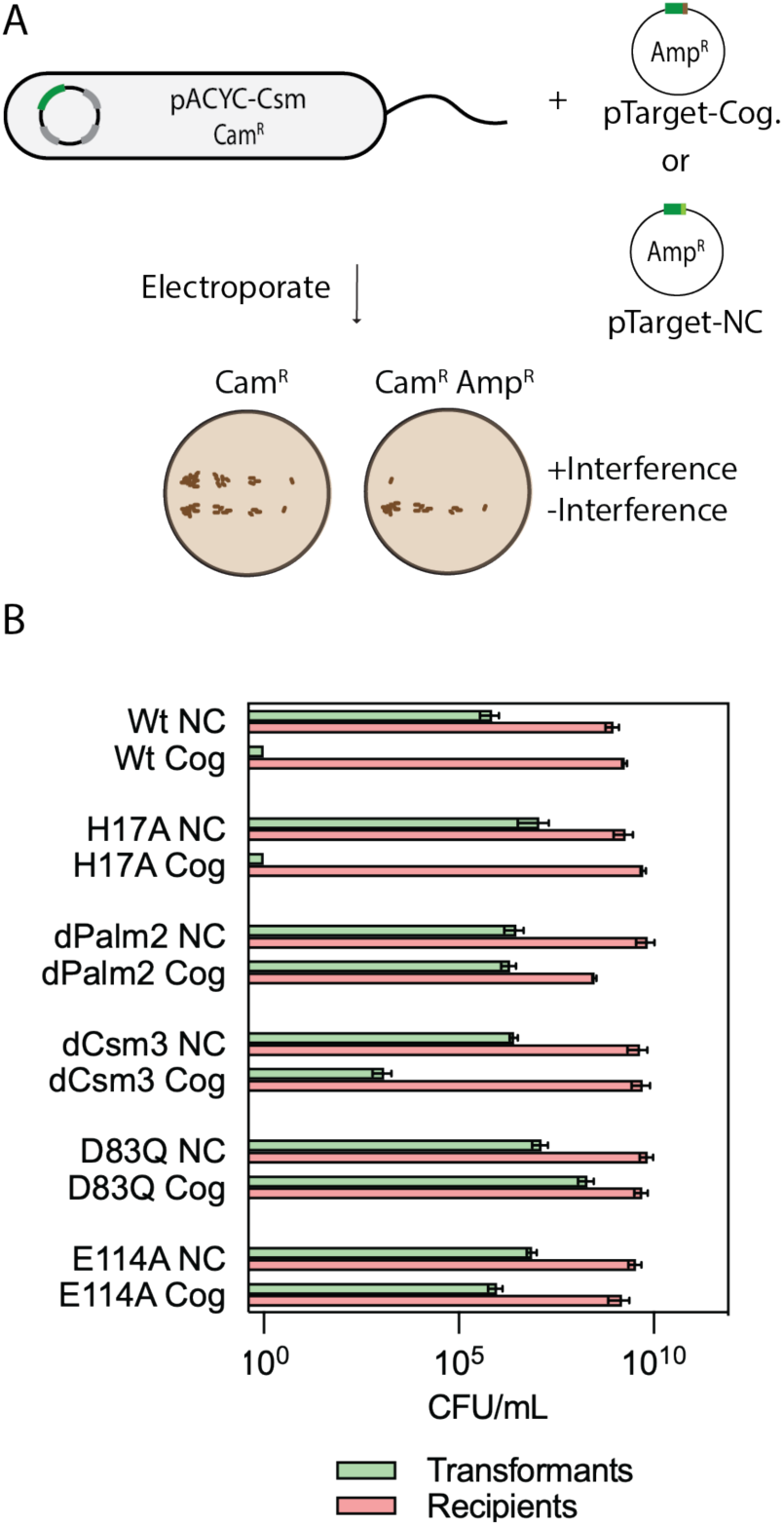
Interference by *S. marcescens* CRISPR-Cas10 is cOA dependent and NucC dependent. (A) Cells carrying pACYC-Csm (recipients) are transformed with pTarget-cognate (Cog) bearing complementarity to crRNA. A pTarget-non-cognate (NC) plasmid is used as a negative control. (B) Site-directed mutants of pACYC-Csm were subjected to the interference assay: H17A (Cas10 HD domain mutant); dPalm2 (Palm2-dead, mutant of GGDD motif); dCsm3 (Csm3-dead, D34A mutant); D83Q (mutant of NucC active site); E114A (mutant of NucC active site). Each bar reports the mean number of colony forming units per mL (CFU/mL) and standard deviation for three distinct transformations. For transformations of wild type with cognate target and H17A with cognate, no colonies were observed. A bar is plotted at 10^0^ to represent these data.

Next, we sought to determine the role of Csm3-mediated target RNA cleavage and cOA activated cleavage of DNA by NucC on interference in our recombinant assay by generating catalytically dead variants for each activity. *S. marcescens* Cas10 is degenerate in its HD nuclease domain and therefore HD domain-mediated DNA cleavage is not expected to occur. Nevertheless, it does contain an H17 residue which we mutated to assay the role of the HD domain on interference. When transformed with pTarget-Cognate, the H17A strain produced no transformants confirming the HD domain does not contribute to interference in our assay (Fig. 3B). An RNase-dead mutant of Csm3 (D34A) produced 10^3^ transformants, an intermediate phenotype for interference, indicating the RNase activity contributes to interference in our assay conditions (Fig. 3B). For mutants defective in cOA synthesis (dPalm2) or in NucC-mediated DNase activity (D83Q or E114A), 10^5^ to 10^6^ transformants are observed in the presence of pTarget-Cognate similar to the transformation efficiency of pTarget-Non-cognate indicating interference has been lost (Fig. 3B). These results show that interference in our system is mediated by cOA-stimulation of NucC DNase activity with Csm3-mediated RNase activity also contributing.

### Characteristics of cOA synthesis by SmCas10-Csm in vitro

Next, we sought to determine the sizes of cOAs synthesized by SmCas10-Csm and assess how mismatches between target and crRNA affect cOA synthesis. To do this, we first purified Cas10-Csm from *E. coli* cells harboring pACYC-Csm. On this plasmid, a hexahistidine tag is appended to the N-terminus of Csm2, allowing the expected 300-400 kDa complex to be purified by immobilized metal affinity chromatography (IMAC) followed by sucrose gradient centrifugation (Fig. 4A). We purified the wild-type complex, a Csm3 D34A mutant complex (dCsm3) lacking RNase activity, and a Cas10 Palm2 mutant complex (dPalm2) defective in cOA synthesis. The dCsm3 complex facilitates characterization of cOA synthesis in vitro, and the dPalm2 complex serves as a negative control. Mass spectrometry confirmed the five prominent bands observed in SDS-PAGE are those of the Cas10-Csm complex (Figs. 4A, Table S3). In the dCsm3 complex the common IMAC contaminant ArnA also appears. ArnA persists in our purification process due to its similar molecular weight in its native hexameric form (445 kDa) (32). We also performed a PAGE analysis of the crRNA that copurified with Cas10-Csm and identified a major band at 42 nt and minor bands at 41 and 48 nt (Fig. 4B). These sizes are consistent with expectations for a type III-A CRISPR complex.

**Figure 4.**
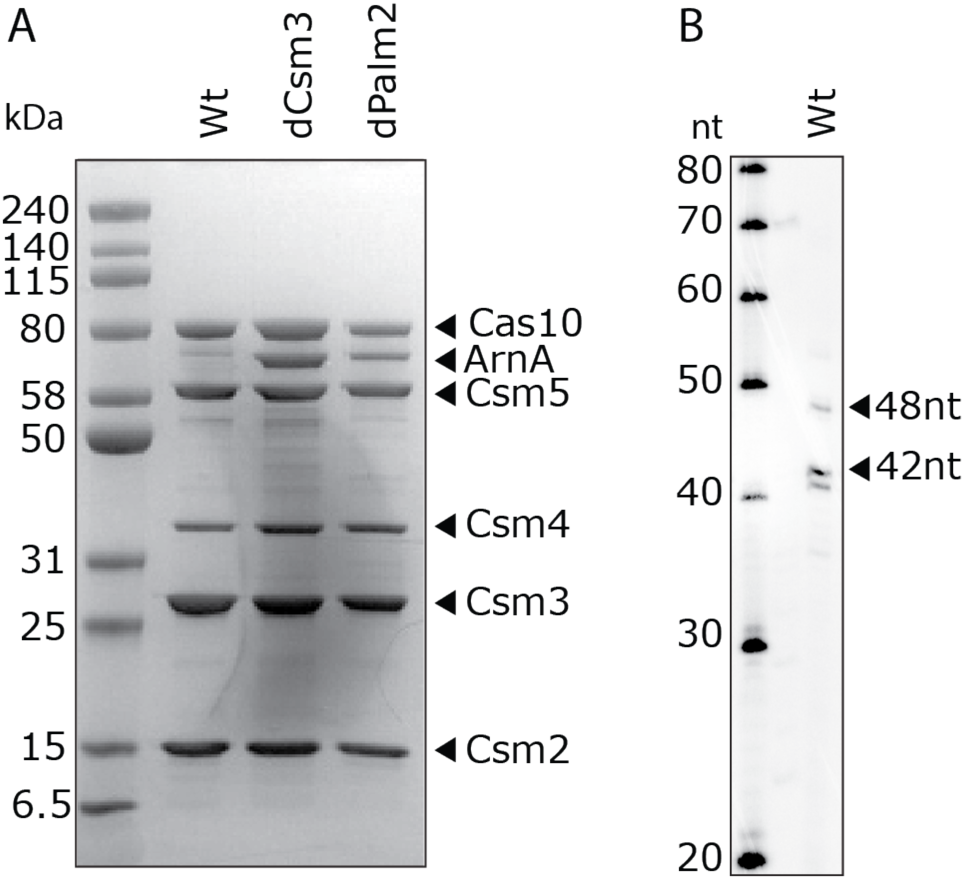
Purification of Cas10-Csm. (A) SDS-PAGE shows that the five proteins that make up Cas10-Csm are present. In the case of Cas10-Csm with a catalytically dead Csm3 (dCsm3) or a catalytically dead Palm2 domain of Cas10 (dPalm2), the ArnA protein, an innocuous common contaminant in IMAC purifications, is also present. (B) PAGE analysis of crRNA extracted from *Serratia* Cas10-Csm. The major species of crRNA is 42 nucleotides (nt) with minor species at 41 and 48 nucleotides.

We sought to measure cOA synthesis by SmCas10-Csm in vitro, including identifying the size of the products. We incubated Cas10-Csm with a cognate target RNA, ATP and Mg^2+^ and subjected the reaction products to MALDI mass spectrometry (Fig. 5A). These data show that cA_3_ (m/z = 987.96) is the major product, with cA_4_ (m/z = 1317.34) present as a minor product. A control reaction with all components except ATP shows that these two peaks disappear (Fig. S3).

**Figure 5.**
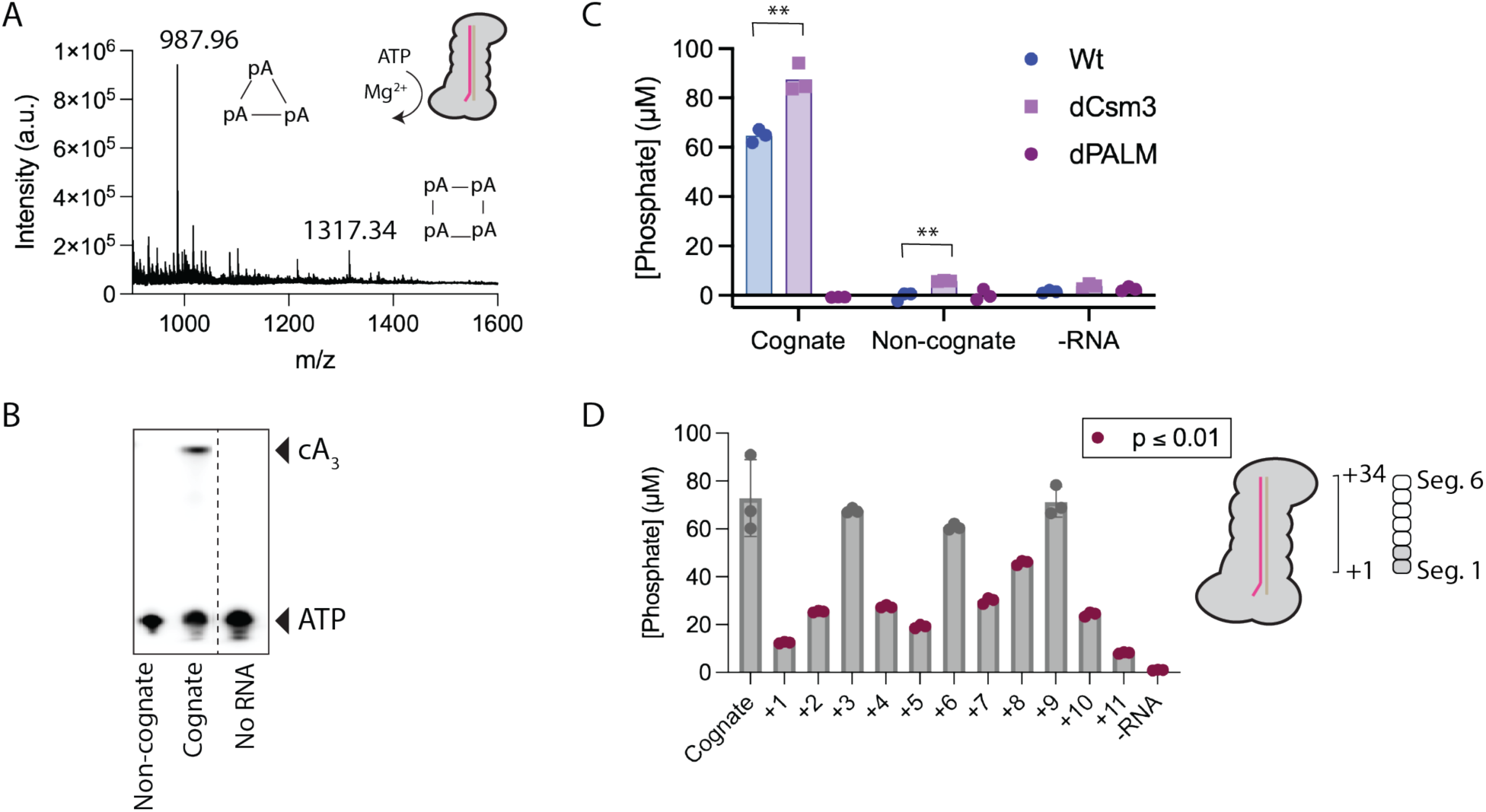
In vitro cOA synthesis by SmCas10-Csm is cognate RNA dependent and affected by segment 1 and segment 2 mismatches. (A) Mass spectrometry provides a clear signal for the presence of cA_3_ at 987.96 m/z (expected mass 988.16 for z = +1) and a modest signal for cA_4_ at 1317.34 (expected mass 1317.21). (B) COA synthesis using α-^32^P-ATP substrate produces a signal on a TLC plate consistent with cA_3_ in the presence of cognate target RNA. (C) COA synthesis by wild type, dCsm3 or dPalm2 complex was assessed by the malachite green assay. Three replicates were performed for each reaction and are plotted along with a bar marking the mean. (D) Mismatches in specific positions of the crRNA-target duplex in segments 1 and 2 reduce cOA synthesis. Three replicates were performed for each reaction. The data is shown as a scatter plot, with bars marking the mean and SEM. The mean of each sample was compared to cognate. Reactions with p ≤ 0.01 are marked in red. ** indicates p < 0.01.

To verify that in vitro synthesis of cOA by SmCas10-Csm was regulated as expected, we performed incubations with synthetic non-cognate target RNA or cognate target RNA and α-^32^P-ATP assaying the reaction by thin-layer chromatography (Fig. 5B). To quantitatively assess the stimulation of cOA synthesis by cognate target RNA we utilized the malachite green assay. We performed experiments with wild type, dCsm3 and dPalm2 complexes. In the presence of cognate target RNA, wild type and dCsm3 readily produced cOA and, as expected, the dPalm2 complex did not. The dCsm3 complex produced slightly more cOA than wild type which can be attributed to its inability to degrade target RNA, which deactivates cOA synthesis (33). In the presence of non-cognate target RNA, cOA levels from wild type and Palm2-dead complex were indistinguishable while the Csm3-dead complex showed a low level of cOA synthesis that was a statistically significant increase over wild type (Fig. 5C). In the absence of target RNA, cOA production was close to baseline (Fig. 5C).

Evidence shows stimulation of cOA synthesis in type III CRISPR systems is sensitive to mismatches proximal to Cas10. The crRNA-target duplex in type III CRISPR systems consists of six-nt segments: five base pairs form, with the nucleotides engaging in stacking interactions, while the sixth nt is unpaired and rotated away from the paired nts. In type III CRISPR systems the two segments proximal to Cas10 compose the Cas10-activating region and base-pairing within this region has an outsized effect on stimulation of cOA synthesis (Fig. 5D) (8,23,33,34). However, the degree to which specific base pairs in the duplex affect cOA synthesis appears to vary from organism to organism and is also affected by the specific target RNA sequence under investigation (34).

We sought to quantitatively determine how mismatches in the Cas10-activating region affect cOA synthesis by *S. marcescens* Cas10-Csm. We used in vitro transcription to generate a panel of target RNAs with a single mismatch systematically introduced into positions +1 to +11 of a cognate target. We combined each of these targets with Cas10-Csm and measured cOA synthesis using the malachite green assay. A reaction without RNA (-RNA) was used as a negative control. Mismatches at positions +1, +2, +4, +5, +7, +8, +10 and +11 were significantly lower (p ≤ 0.01) than that of cognate target (Fig. 5D). Positions +1 and +11 had the largest defects. The mismatch at +1 led to a 5.8-fold reduction in cOA synthesized.

### The structure of *S. marcescens* Cas10-Csm unbound and bound to target RNA

Conformational change upon binding to target RNA has been observed for several type III CRISPR systems and it is believed that these changes allosterically activate cOA synthesis by Cas10 (23,35–37). However, there is no agreement regarding the detailed mechanism of the allosteric activation. To shed additional light on the structural basis for target RNA mediated activation of cOA synthesis, we used cryo-EM to determine structures of Cas10-Csm unbound and bound to target RNA (Figs. 6A, 6B, S4 and Table S4). We used dCsm3 complex so that our target bound structure would possess intact, rather than cleaved target RNA. The Cas10-Csm structure without target RNA was determined to 4.48 Å resolution (FSC_0.143_). AlphaFold models of the Cas10-Csm proteins were rigid-body fitted into this map, indicating the presence of the Cas10, Csm4, Csm3 and Csm5 proteins (Fig. 6A). Csm4 is bound to the 5’ tag of crRNA and also forms protein-protein interactions with Cas10. Four copies of Csm3 form a filament gripping bound crRNA and presenting it to solution. Csm5 binds the 3’ end of crRNA and caps the Csm3 filament (Fig. 6A). Little density is seen for Csm2, despite the fact that it is clearly present in the complex by SDS-PAGE (Figs. 4A, 6A). This indicates Csm2 is highly dynamic in the absence of target RNA. Domain 4 of Cas10 which interacts with Csm2 is also partially disordered in unbound Cas10-Csm (Fig. 6A).

**Figure 6.**
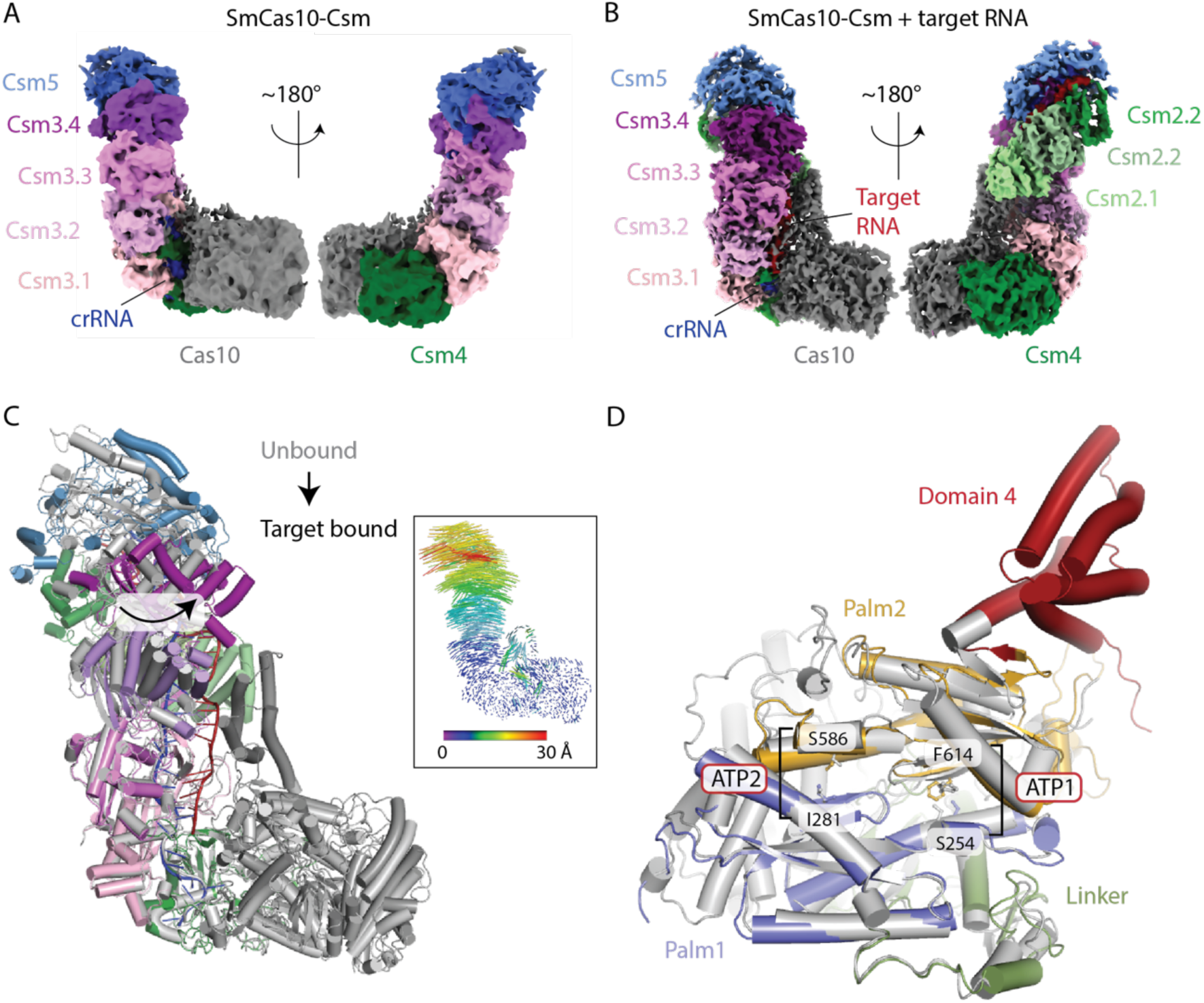
Conformational changes upon target RNA binding to SmCas10-Csm. (A) The cryo-EM map of SmCas10-Csm in the absence of target RNA. CrRNA and four of the five Csm proteins are visible. Four copies of Csm3 (Csm3.1-Csm3.4) are present. Csm2 is too dynamic in the absence of target RNA to be resolved and little density is present for domain 4 of Cas10. (B) In the cryo-EM map of SmCas10-Csm bound to target RNA, Csm2 is resolved, as well as domain 4 of Cas10. (C) A superposition of unbound versus target bound SmCas10-Csm reveals conformational changes throughout the complex. Large rotations of the distal end of the complex (Csm5 and Csm3.4) occur. A vector depiction of these changes is shown as an inset. (D) A view of conformational changes in Cas10 induced by target RNA binding which include ordering of domain 4. The cOA synthesis active site, which resides at the interface of Palm1 and Palm2 is marked by the residues that mediate ATP binding at the ATP1 and ATP2 sites. Re-organization of the ATP1 site is associated with target RNA binding. Unbound Cas10 is shown in grey and bound Cas10 in color

We also determined the structure of the target-bound Cas10-Csm complex to a resolution of 3.84Å (FSC_0.143_). In this structure, Csm2 and domain 4 of Cas10 are clearly resolved in addition to Cas10, Csm4, Csm3 and Csm5 (Fig. 6B).When the two structures were superimposed (Fig. 6C), the Csm3 filament and Csm5 can be seen to rotate toward the minor groove of the crRNA-target duplex by as much as 30Å (Fig. 6C). Large movements of the now well ordered domain 4 of Cas10 are also observed. The active site for cOA synthesis lies in a composite surface between the Palm1 and Palm2 domains with residues from both domains contributing to ATP binding (37–39). Domain 4 of Cas10 extensively interacts with Palm2 suggesting that movement of domain 4 should affect the position of Palm2. Indeed, we see changes in the position of the β-strand which contains F614 and the alpha-helix that contains S254, two residues that form the ATP1 site, one of the two ATP-binding sites required for cOA synthesis (Fig. 6D). In contrast, the ATP2 site, indicated by sequence homology to be formed by residues I281 and S586, is minimally changed by target RNA binding. The structural data suggest that conformational changes in domain 4 of Cas10, which in turn influence the organization of the ATP1 site, may underpin the allosteric mechanism for activation of cOA synthesis by target RNA binding.

Close inspection of the region in which domain 4 contacts Palm2 reveals a network of interactions, also involving Csm3, that form upon target binding. A β-hairpin in Palm2 encompassing residues Y612-G624 contains the conserved active site GGDD motif (G616-D619), which binds a Mg^2+^ ion that is critical for catalysis (Fig. 7A). The aforementioned F614 that participates in the ATP1 site is part of this β-hairpin. The conserved W626 residue is located just following the C-terminus of the β-hairpin (Fig. 7A). Interestingly, in the target-unbound structure W626 of the Palm2 domain and Y807 of domain 4 and the Csm3 residue I30 are all far from each other (Fig. 7B). However, upon target binding movements of Csm3 loop containing I30 and domain 4 bring each of these residues, along with the Csm3 residue L28 into proximity and van der Waals interactions form among them (Fig. 7C). It is likely that the van der Waals network that forms between residues in Csm3, domain 4 and Palm2 plays a key role in organizing the ATP1 site for catalysis upon binding of target RNA to Cas10-Csm (Fig. 8).

**Figure 7.**
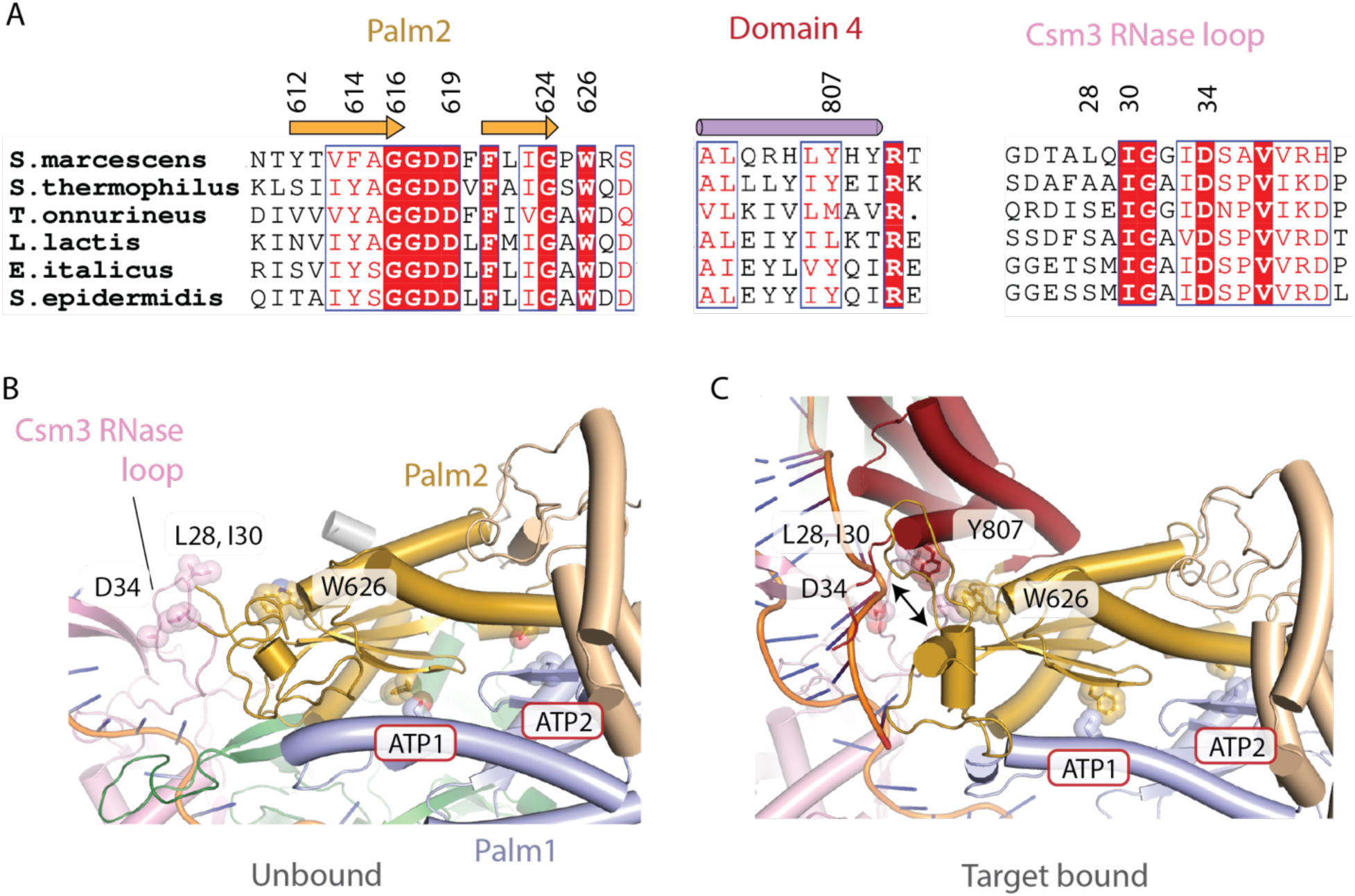
A network of van der Waals interactions adjacent to the ATP1 site is altered on target RNA binding. (A) A structure-based sequence alignment of Palm2 and domain 4 was constructed for Cas10 from six prominent model systems. W626 is conserved and Y807 is conserved for four out of six species. A Csm3 sequence alignment shows variation at L28 but strict conservation for I30. D34 is the catalytic residue for the Csm3 RNase site. (B) A gap exists between Palm2 residue W626 and Csm3 residue I30. (C) However, upon target binding, a network of van der Waals interactions forms between W626, domain 4 residue Y807 and Csm3 residues L28 and I30 that may facilitate formation of an ATP1 site active in cOA synthesis.

**Figure 8.**
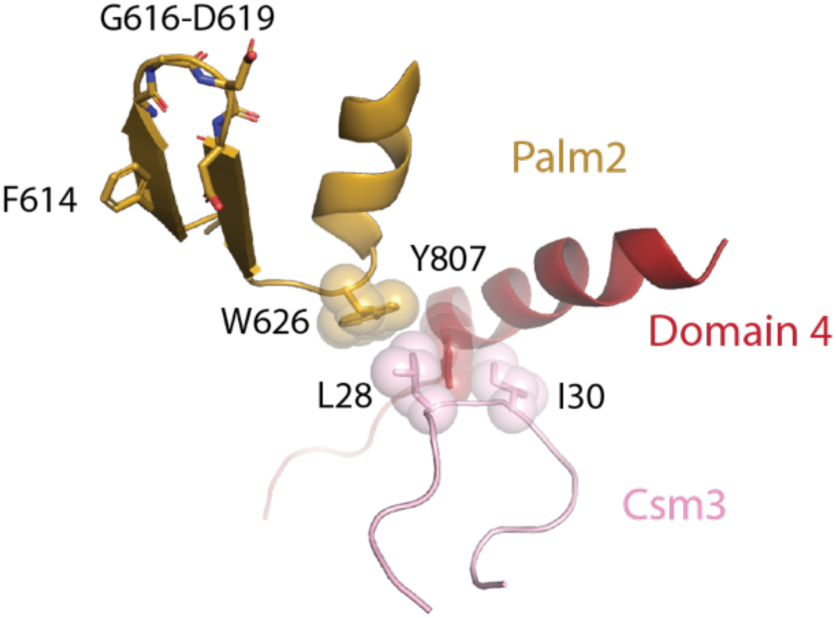
A network of van der Waals interactions among Csm3, domain 4 and Palm2 that forms upon target RNA binding.

### SNP detection in RNAs by Cas10-Csm

Sickle cell disease (SCD) is caused by mutations in the human β-globin gene (*HBB).* Since SCD poses a major burden in sub-Saharan Africa where there are large populations without access to advanced diagnostics, we asked whether the *HBB* transcript and its most common variant could be detected by Cas10-Csm programmed with a complementary crRNA (40). To date, Cas10-Csm has not been used for SNP identification on human transcripts, but since cOA synthesis is sensitive to single-nucleotide mismatches in at least some positions of the crRNA-target duplex, we reasoned that it could be done (Figure 5D). The most common SCD-associated allele possesses a SNP, G<u>A</u>G to G<u>T</u>G, leading to a Glu to Val change in the sixth amino acid of the mature human β-globin protein [NM_000518.5:c.20A>T (p.Glu6Val)] widely referred to as the HbS allele. We will denote the corresponding transcript as A20T from here on. We sought to detect the wild type *HBB* transcript and the A20T transcript with Cas10-Csm programmed with a complementary crRNA. We designed our Cas10-Csm expression construct to encode a crRNA that would base-pair to the *HBB* transcript such that the SNP would be located at the +1 position (Fig. 9A). For read-out, we established a fluorescence-based reporter assay in which *HBB* transcript detection by Cas10-Csm stimulates cOA synthesis, which in turn stimulates NucC to cleave a synthetic DNA duplex labeled with a fluorophore-quencher pair liberating fluorescence (Fig. 9B). CrRNA-B01 is designed to recognize wild type *HBB* and crRNA-S01 is designed to recognize A20T.

**Figure 9.**
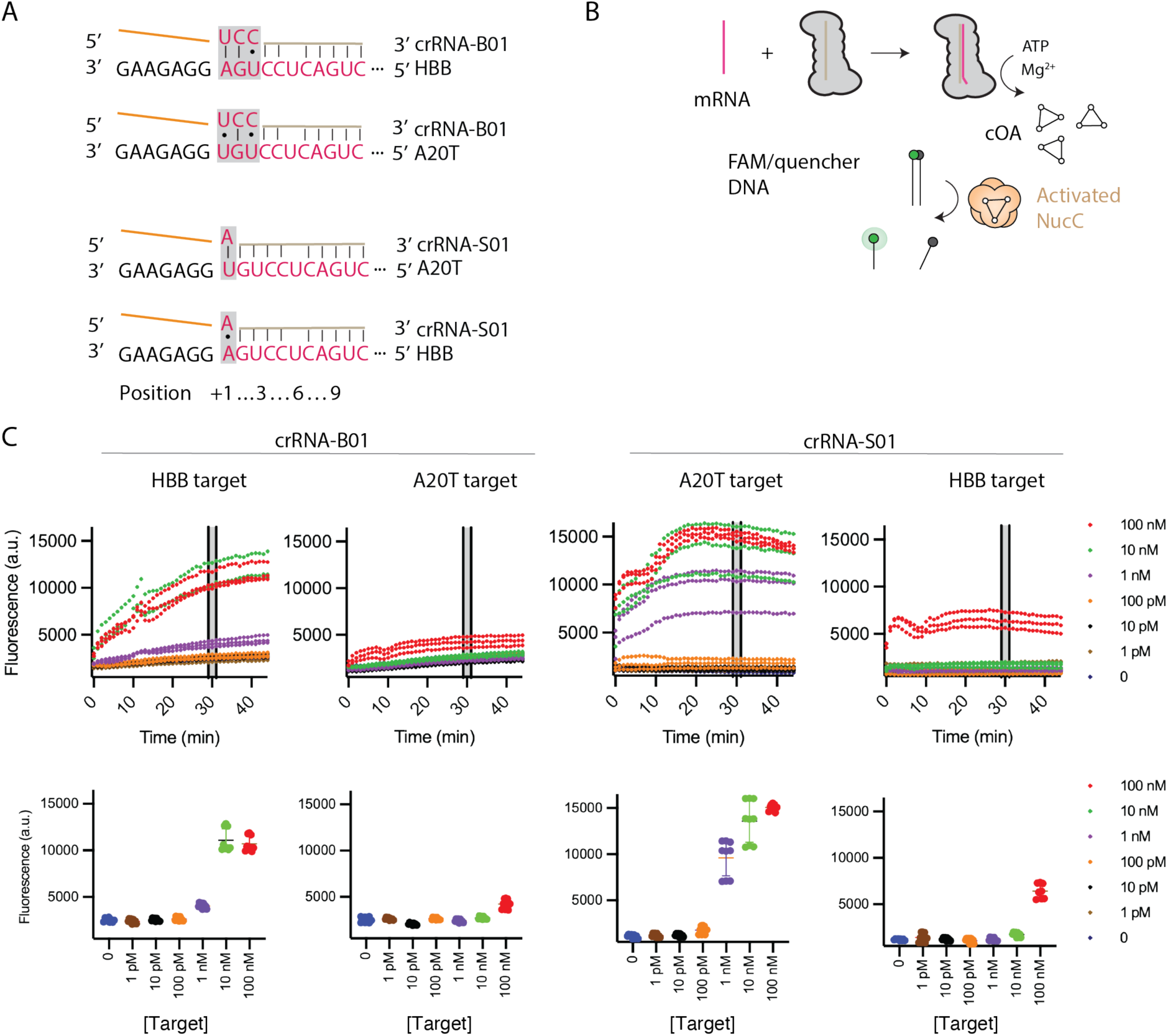
Detection of human β-globin RNA by Cas10-Csm in a NucC coupled assay. (A) Base-pairing between the crRNA probe and the target RNA at position +1 is used to discriminate wild type *HBB* sequence from the disease causing *HBB* c.A20>T (A20T) sequence. CrRNA-B01 was designed with a pyrimidine-pyrimidine mismatch at +3, which enhances the ability of this crRNA probe to discriminate between the sequences. (B) When Cas10-Csm is incubated with a RNA complementary to its bound crRNA, cOA is produced, the DNase activity of NucC is stimulated and cleavage of a reporter DNA occurs. (C) In the upper plots, Cas10-Csm bearing a crRNA designed to sense an HBB or A20T sequence was subjected to our fluorogenic assay with either a HBB target RNA or an A20T target RNA. In the lower plots, fluorescence values from the timepoints at 29, 30 and 31 minutes from three replicate reactions were plotted versus target RNA concentration.

We incubated Cas10-Csm bound to crRNA-B01 with RNA mimicking either wild type *HBB* transcript or A20T transcript for 2.5 minutes to generate cOA and then added our read-out mixture including FAM/quencher labeled DNA and NucC (Fig. 9B). We performed reactions with titrated target RNA and used the fluorescence signal from the 29-31 minute timepoints to detection events. We chose an I/σ(I) of 3.0 for calling detection. The reaction with crRNA-B01 and wild type *HBB* RNA exceeded the detection threshold at 1 nM. When this complex was presented with A20T RNA, the fluorescence signal at our detection threshold was only observed at 100 nM A20T (Fig. 9C). This suggests weak cOA stimulation of crRNA-B01 complex by A20T and indicates a broad two-orders-of-magnitude window in which the complex can readily discriminate between *HBB* RNA and A20T RNA (Table S5). The reticulocyte contents of human blood contain abundant *HBB* transcript (nM levels in whole blood), so transcript captured from several mL of human blood should be easily detectable by our fluorescence assay. We also incubated Cas10-Csm bound to crRNA-S01 with titrations of A20T RNA and wild type *HBB* RNA. This Cas10-Csm complex also achieved detection for A20T RNA at 1 nM concentration but 100 nM of wild type *HBB* RNA was required to produce substantial fluorescence, again revealing a two orders-of-magnitude window in which specific detection can be achieved (Fig. 9C and Table S5).

To further test the idea that Cas10-Csm could have use in distinguishing wild type *HBB* transcripts from A20T transcripts in RNA extracted from human blood, we constructed contrived samples. To total human RNA from liver cells, which do not express β-globin, we added either synthetic A20T RNA, *HBB* RNA or both to simulate three genotypes that would commonly be encountered: homozygous wild type, heterozygous or homozygous for A20T. Fluorescence versus time curves were collected and data from the 29-31 minute mark were compared among the samples. CrRNA-B01 complex produced a large fluorescence signal in the presence of 5 nM *HBB* RNA or a mixture of 2.5 nM *HBB* RNA and 2.5 nM A20T (Fig. 10A). A low level of fluorescence was observed when crRNA-B01 complex was incubated with 5 nM of A20T RNA, however, a t-test (two-tailed) demonstrated the samples could be distinguished from each other with p < 0.0001. CrRNA-S01 complex was also incubated with 5 nM *HBB* RNA, 2.5 nM *HBB* RNA and 2.5 nM A20T or 5 nM of A20T RNA. An analogous pattern was observed, a strong fluorescence signal was observed when A20T was present, but the crRNA-S01 complex produced only a weak signal similar to our no RNA control when given wild type *HBB* RNA. A t-test demonstrated A20T containing RNA can be distinguished from wild type *HBB* RNA with p < 0.0001. In sum, these proof-of-concept data indicate that the use of Cas10-Csm for SNP typing merits further exploration.

**Figure 10.**
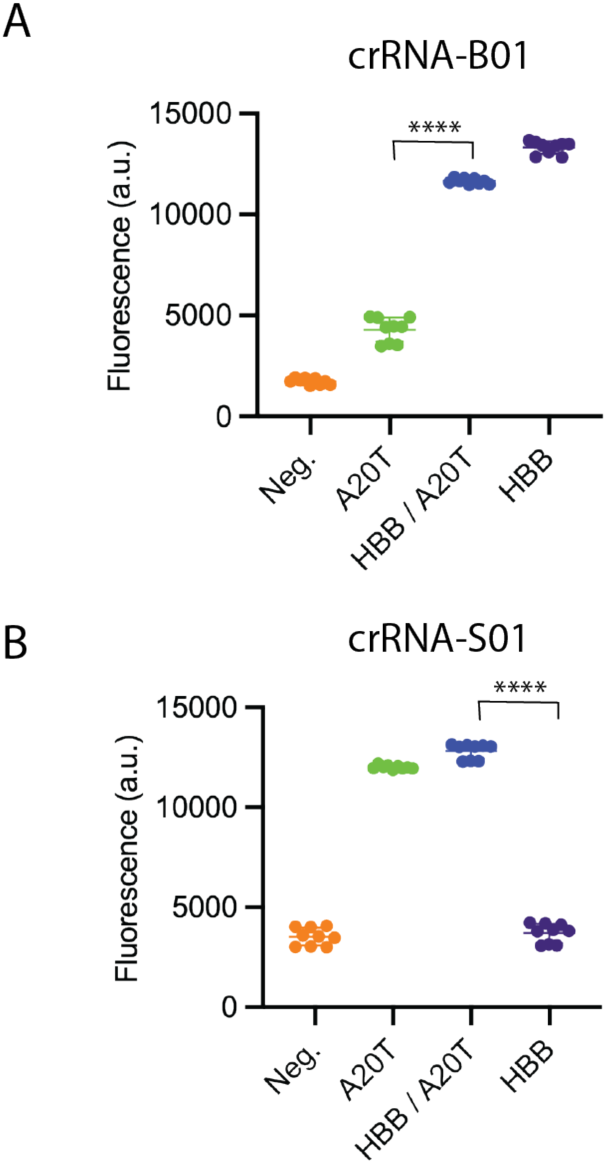
Cas10-Csm can distinguish between wild type and disease-associated hemoglobin-β RNA in contrived samples. (A) A contrived sample was constructed by adding 5 nM wild type HBB RNA, 2.5 nM HBB RNA and 2.5 nM A20T RNA or 5 nM A20T RNA to human liver total RNA. The mixture was probed with Cas10-Csm bound to crRNA-B01 and subjected to our fluorogenic assay. (B) The same assay was performed as in (A) except Cas10-Csm was bound to crRNA-S01. A two-tailed T-test was performed. ****, p < 0.0001

## Discussion

We established a recombinant interference assay for SmCas10-Csm and demonstrated that interference is dependent on cOA synthesis and NucC DNase activity. We established a recombinant expression and purification procedure for SmCas10-Csm, demonstrated its ability to synthesize cOA in vitro, assayed the sensitivity of this synthesis to mismatches with target RNA and determined the cryo-EM structure of SmCas10-Csm unbound and bound to target RNA. The structures shed light on the conformational dynamics associated with Cas10 activation while the cOA synthesis assays suggest that a large enough difference in cOA production exists when stimulated by cognate target versus RNA with a single mismatch to make SNP detection possible. We tested this idea by programming SmCas10-Csm with a crRNA complementary to RNAs mimicking the human hemoglobin β transcripts and were able to distinguish between wild type and a disease-associated variant.

Many intricate systems have recently been uncovered that allow bacteria to defend against phage infection, but phages in turn possess novel mechanisms to counteract these defenses (13,14,41). Recently it was discovered that jumbo phages can encapsulate their genome inside a proteinaceous phage nucleus during bacterial infection to protect it from attack by restriction-modification systems and DNA targeting CRISPR systems, such as type I or type II systems (16,17,42). Jumbo phage transcripts, however, must leave the phage nucleus to be translated and thereby become detectable by type III CRISPR systems, such as Cas10-Csm. An important model system for studying this phenomenon has emerged. Utilizing infection of *S. marcescens* by the jumbo phage PCH45, it has been shown that the Cas10-Csm of *S. marcescens* can detect phage transcripts, leading to cA_3_ synthesis by Cas10-Csm (11). The cA_3_ activates the DNase activity of NucC, which shreds bacterial genomic DNA, blocking phage replication through an abortive infection mechanism (11). Our results demonstrating the reconstitution of SmCas10-Csm activity in vitro, its sensitivity to mismatches in target RNA, and its structure will facilitate future development of the *S. marcescens* model system for studying the dynamics of jumbo phage infection and the cell’s Cas10-Csm mediated response.

Regulation of Cas10 activity is dependent on two features of target RNA: sufficient base-pairing between the body of the target and crRNA and the presence of the unpaired 3’ flank of the target (34,43,44). The unpaired 3’ flank appears to be sensed by a direct interaction with the junctions of the Palm1-linker and Palm2 domains of Cas10 that has been observed in several structures (35–37). This interaction is not captured, however, in the majority of structures for type III complexes, suggesting its transience (45–48). All structures of type III complexes indicate that a target RNA body region base-paired to crRNA is sensed by interactions with domain 4 of Cas10 and through interactions with the Cas11 homolog of the complex (Csm2 or Cmr5). However, comparison across multiple structures again indicates a diverging picture of the complex’s dynamics. In structures of the *Lactococcus lactis* Cas10-Csm complex, density for Csm2 is only clearly observed in cryo-EM maps once target RNA is bound indicating the dynamic nature of the protein in the unbound complex (36). We observed the same phenomenon for Csm2 in our SmCas10-Csm structures suggesting this behavior of Csm2 is not unique to *L. lactis*. Additionally, our cryo-EM maps indicated a disordered-ordered transition for domain 4 of Cas10 upon target RNA binding that had not previously been observed. Detailed descriptions are emerging for how the rearrangements associated with target RNA binding activate Cas10 for cOA synthesis and HD-domain mediated DNA cleavage (37,49). Specific contacts between domain 4 of Cas10 and target RNA are required for cOA synthesis (50). We have offered, based on the cryo-EM structures we report, a model that proposes re-ordering of the domain 4-Palm2 interface for cOA synthesis is influenced by a structural nexus that forms between Cas10 domains 4 and Palm2 and Csm3.

The earliest demonstrations of CRISPR systems as point-of-care molecular diagnostics used Cas9, Cas12 and Cas13 (18,20,51,52). These systems display good specificity, but in most cases rely on amplification of the target nucleic acid before CRISPR-based detection to achieve high specificity, which adds complexity to the assay (21). Recently, Cas10-based systems have been used as nucleic acid diagnostics (8,22–24,53). These systems take advantage of the intrinsic signal amplification associated with Cas10: a single RNA detection event activates multi-turnover cOA production by Cas10 and these cOA activate a multi-turnover reaction in the Csm6 (Crf1) or NucC effector. Cas10-based systems have the additional benefit that cOA can stimulate a diverse family of enzymes giving flexibility to the choice of read-out chemistry. An outstanding question, however, is the specificity of Cas10-based molecular diagnostics given that type III systems have been shown in many cases to be promiscuous in vivo (26,34,54–56). We demonstrate the ability of Cas10-Csm to distinguish two variants of the human *HBB* RNA, the wild-type and the variant with the SNP that causes the Glu6Val mutation, which is the most common variant associated with SCD (57). CRISPR-based diagnostics like the one we report could be useful for genotyping individuals from a few mL of blood in the low resource health care settings that exist in many regions of Africa where SCD is widespread.

## Data availability

Proteomics data are available at ProteomeXchange with the identifier PXD074731. The coordinates describing Cas10-Csm unbound to target RNA and bound to target RNA are deposited in the Protein Data Bank with deposition codes 9zs2 and 9zs4. The corresponding maps are deposited with the Electron Microscopy Data Bank as EMD-74675 and EMD-74688.

## Funding

Research reported in this publication was supported by the National Institute of General Medical Sciences of the National Institutes of Health under Award Number R35GM142966 to J.A.D.

## Conflicts of interest statement

The authors report no conflict of interest.

## Acknowledgements

Electron microscopy was carried out at the University of Alabama at Birmingham (UAB) Cryo-EM Facility (CEMF), funded by the UAB Institutional Research Core Program, the O’Neal Comprehensive Cancer Center (NIH grant P30 CA013148) and National Institutes of Health grant S10 OD024978 to T.D.

## Notes

### Competing Interest Statement

The authors have declared no competing interest.

