## Supporting Information for "Structural insights into target detection by the *S. marcescens* type III CRISPR complex and its deployment in SNP identification"

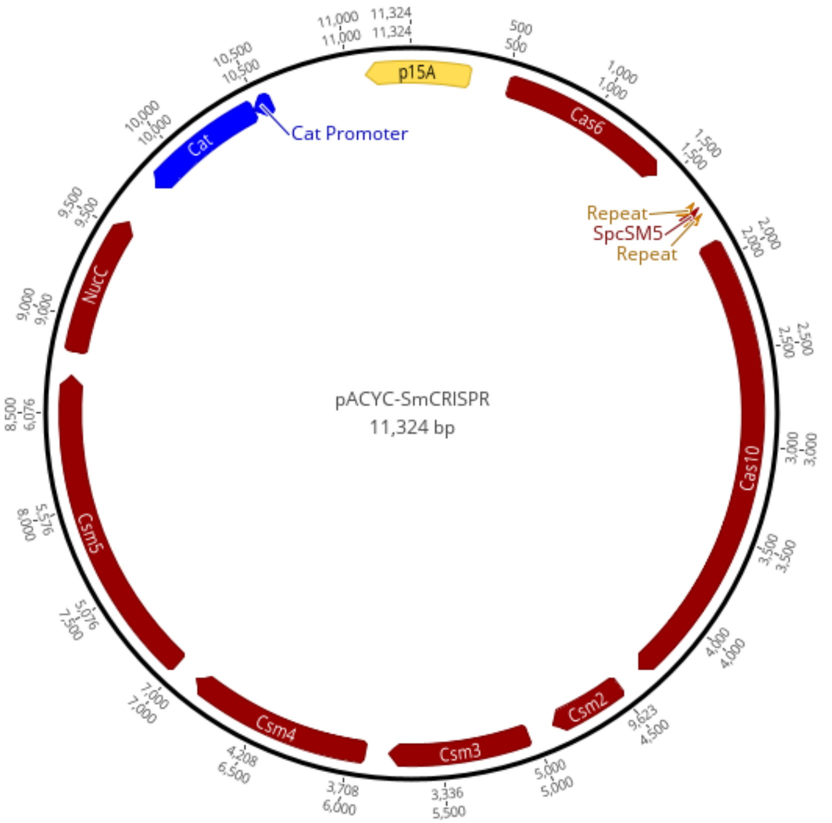


**Figure S1. Plasmid map for pACYC-Csm**

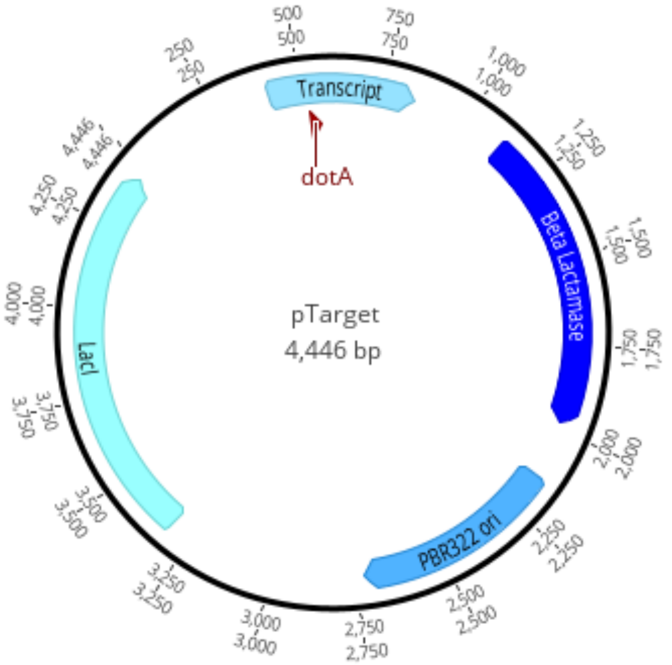


**Figure S2. Plasmid map for pTarget**

**Table S1. Plasmid sequences, gene fragments and oligos used in the study.**

| Name | Sequence (5'-3') |
|---|---|
| Plasmids | |
| pACYC-Csm | 1 atgcacgaac cccccgttca gtccgaccgc tgcgccttat ccggtaacta tcgtcttgag<br>61 tccaacccgg aaagacatgc aaaagcacca ctggcagcag ccactggtaa ttgatttaga<br>121 ggagttagtc ttgaagtcat gcgccggtta aggctaaact gaaaggacaa gttttggtga<br>181 ctgcgctcct ccaagccagt tacctcggtt caaagagttg gtagctcaga gaaccttcga<br>241 aaaaccgccc tgcaaggcgg ttttttcgtt ttcagagcaa gagattacgc gcagaccaaa<br>301 acgatctcaa gaagatcatc ttattaatca gataaaatat ttctagattt cagtgcaatt<br>361 tatctcttca aatgtagcac ctgaagtcag ccccatacga tataagttgt aattctcatg<br>421 tttgacagcc ttcagatccg atagactagc cgctggtaat aatacgactc actataggga<br>481 gagaattcta agaccgaaag tcggaaacaa agaggattta tatgacatgg gtttgccatc<br>541 tggcgcgtta ccgcctctgt tttcaaagca tcgaaccgct ggcgctgccg ctgtttagcg<br>601 gctccatgct gcgcggcgcg ttcggccatg cgctgcggcg catctgttgc atcagccgtc<br>661 agaaacagtg tgacggctgc ccgttgctgg ccggttgcca gtacccgctg ttgtttgagc<br>721 cgcgcttgct gcaaaacacc cccggccagc cccaaccggc tccgccttat gtgctggagc<br>781 cagcggccgt tcagagggag cttgccgcag gggaaatctg gccggtagac gtggtgctgc<br>841 acggcccggc gctaccgcac ctgagcctga ttattctggc ctggatgcag gcagctgttc<br>901 aggggtttgg cagccagcgg gtgccagcac agttgtgcca ggtgcaggta gagcaacccg<br>961 atactgaagc gcgctggctg acgatctggc gtcacgacca gccgtttatc ctgccacacg<br>1021 ccaccacgca aacaccgcca ccgcccccgg cggcagaggc gctgcgcctg cacttcacca<br>1081 ccccgacccg gctgttgcgc caggggcaac tggtacgggt gcgtgaactt caggcacacg<br>1141 acctgctggc cgcgctggag cgacgcctgc acacgctggc cgcctcgctg ggcattgccc<br>1201 cgccgccgtc gctgaccacg caactcgtta cgctgcaacc cggccagttg cgctggatga<br>1261 actggcaacg ctactccagc cgccagcaac aggagatgaa tctcggcgga tttatcggtg<br>1321 acgttacgct ccacggcgag ctgacgccgc tgtggacatg gctgtggctg ggccagtggt<br>1381 tgcatgtggg caaaaacagc agcttcgggc tgggacgcta tcagttagag acggttcccc<br>1441 gctaaaagct ttcctgtgag cagcgaaagc ctagcataac cccttggggc ctctaaacgg<br>1501 gtcttgaggg gttttttgtt atacgcgaga taatcacttg catagctgcg tatggaggaa<br>1561 gcaactcttg agtgttaata tgttgacccc tgtattaggg atgcgggtag tagatgtggg<br>1621 cagagacacc cacactgcca gatcttaata cgactcacta tagggagacc atgggtcctt<br>1681 acggacgctc cctgactgaa gggattaaga ccatttacag aacctgttcc acttaatgaa<br>1741 gttgcgtcct tacggacgct ccctgactga agggattaag acctcgaggc tgtggtctag<br>1801 acattccata catatcgggg gggtaggggt tttttgtgtg cctctagtgg ctggctaaga<br>1861 ataatacgac tcactatagg gagaggatcc ataaaggagg taaataatga actggcttgc<br>1921 cgcctcttgc catgtggctg cctttgcgct gttacataac ctgaaaccgc tggcgcagcg<br>1981 tgccggcatt caggatttcc cgtcaccggc gctggcaaac aaactgtttg cgccgctgcc<br>2041 ggaaagcctc tcggcccgtg cggatctgca cgaaaataca ttaaccctgc tattggattt<br>2101 cgccgccaga ctggcgcgcg gcctgccgga gacgcccgta agccgtggcg aaaccgggct<br>2161 gataccgctc acccacctgt tagcggacga taaccacagc gccccggcgg agcgctgtta<br>2221 cagcccactg gccccgctgg gggtcgattc gctgatgccg gtgacggcag cgttgcccac<br>2281 cggcgaacgg caggcggcgt accggcaggt gtatcaggca ttgatagagg ggctggaggc<br>2341 gatccccgcc gcccaccgcc tgcaaccgtc gctgtggctt gaccacctcg acagcctgtg<br>2401 gatgaccacc tgccacgcgc tgccagacgg cgacggggcc agcggtatct cgctgtacga<br>2461 tcagggcaaa accacggcgg cgctcgccgt cgcactgtgg cagcaccacg cccgtcaggc<br>2521 cacaccggag aaggaaattt cactcacgcc aggcgacgat gcgcgcctgc tgctgattca<br>2581 ggccgatgtg ttcggcattc aagagctgat tttcgcccag ggcaatcaga cgcaaaaaat<br>2641 ggccgccaaa ctgctgcgcg ggcgttcgtt tcaggtatcc ctgctggcgg aaaccgctgc<br>2701 gctgcgggtg ctggaaacct ttgacctgcc gccggtttgc cagttgatta acgccgccgg<br>2761 caaaagcctg attgtcgcgc cgaatctgcc ggatgccgcc gaacggctgg ccgggttgcg<br>2821 tcagcgtctt gatgagtggt ttttaaccca cacctacgcg caaaccggca tcgggctgtg<br>2881 cagcacggtg gccggcagcg ctgagttcct cggcgagcag gcttacggca ggctgcaaaa<br>2941 ccggctggcg caggcgatgg agcaacagaa ataccagcgt tttttccctgt gtgccggtga<br>3001 tgcgcccccg ccggtgtttg agggttatct ggaccaaata gcgcagggcg caggcggtga<br>3061 accctgcccg atgaatggcc tgcacccggt ggaaaccacc ctcaacgggc ttggctgtag<br>3121 ccgactggcg gcagaccaga taaccctcgg ccactggata aaccacggcc actggataag<br>3181 ccacggcgag cgttacgccc ggctgttgat cctgcgtgac agcaccgcca ccttctctga<br>3241 tgaccgctgc ctgcacctga cgctgtttgg ctatcaggtg gtggccgtca gcctgaaaga<br>3301 ggacagcggc gagttcggcg aactggcgtg caacggcagc ctgcgccgct gctgggatgt<br>3361 cagcctgccg ccggaaggca cctccctgtt tcagggttat gcccgccgtt ttatcaacgg<br>3421 ctggatgccg ctggcgcagg gggaatatca gccggagtgc taccccggca ttgaagaagc<br>3481 gctcgctaac ggcgacatca aaacctttga tcacctgagc tgtgaagatt tgtatcagga<br>3541 tgccgaccag tcgttacgcg gcacctgcgc gctgggcgtg ctgaaagggg atatcgacaa<br>3601 cctcggccac ctgtttcgca gcggcctgcc gcaaccgggc tttgccaaaa ccatcggcct |

3661 gtcgcgccag atacacctgt ttttcactct gtggctgccg catctgtgcc gcaaagaccc
3721 gcgttttgcc aacacctaca ccgtgttcgc cggcggcgac gatttctttt tgattggccc
3781 gtggcgcagc cagcaaacgc tggcgctaac aatggcgcag gacttcgcgc gctacagcgg
3841 ccacaatccg gcgctgcatt tttcactcgg gctggtgcag gccaaacccg gctacccggt
3901 gcgggcgctg gcggctcagg ccgaagccgc tctgaagcag gccaagcaac accccggcaa
3961 gaacgccatc tgcctgtata acgaggtgat gggctggccg gaatatgacg cgttactggc
4021 gtgcagtgag gagctggcgc gctggcgcga gcatgacggc tacccgctct cctccggcct
4081 gctttaccgg ttgctggcgc tcagtgaaca gtcggcggag gagtcagaaa aaccgcaggc
4141 ggcactgtgg cgcagccggc tggcctattt tttacgtcgc aatctggtgg ataacgtgaa
4201 agtcccgacg aaagaaaacc cggcagcgtt ccgccatcag cttcatttgg aattgtttga
4261 gaaacttgag cagcacctga aacggcaccg ccagcgttac cgggtggcgc ttcaacgcca
4321 tctctaccac taccgcacgg tgccgcgcgg cgagtaacat atggctgcgt ggtcaaatgt
4381 gcgtacccta accccttccc cggtcaatcg ggcggatgg ggtttttgt gcgtacttca
4441 ttatgtatat taatacgact cactataggg agaagatcta taaaggaggt aaataatgca
4501 tcaccatcac catcacacga cgctcgacta ttttaaacgc caactgcaca agccggaggc
4561 gacatttttt gatgacgacg cccaaaaatt tgccgaacag ttggctcaat acggtaagaa
4621 gggcaaaccc actcagttgc gccgctttta cgaccagtta cagcaactgg agcagcgcat
4681 caatggcgat gaggaaaaac tggctctcta cctacccgag atccgcatga tcagcgccca
4741 tctggcgtat gccaaagggc gcgagcttat ctctgacgag ttttgccaaa ccatgcaaaa
4801 cctgattcgc agcatcaaca cctgccagca cctgaaaaat ggccggctgt tttttgaagc
4861 caccctcggt tttttacgcg ccatacgccg cgactgaacg cgtgctgcgt ggtcaaatgt
4921 gcgtagacca accccttgcg gcctcaatcg ggggggatgg ggtttttttgt caggcaagtc
4981 tcagctggtt taatacgact cactataggg agagaattca taaaggaggt aaataatgca
5041 actgaacaat atccagacgt tacgcgccac tctggtgtgt gaaaccgggt tacacattgg
5101 cggcggcgac accgcgttgc agattggcgg catcgacagc gccgtggtgc gccacccgtt
5161 gacccagcaa ccttacattc ccggctccag cctgaaaggc aaactgcgca gcctgctcga
5221 atggcgcgct ggtgtggtcg gcgataccga gggcaaggtg ctcagtcatc aggtttacca
5281 gcaactgatc gacgataaaa aacaggcaca ggcactacag gcactgaaaa tcctgcaact
5341 gtttggcgtc agcggcggcg acaagctttc cgccgaacaa gcccaacaga tcggcccgac
5401 tcgcctgtcc ttctgggatt gcgaatttga cgaacactgg ctggcgcagc aaggcgggcg
5461 cgtccagacc gaagagaaag cggaaaactg tattgaccgc atcagtggcg tggcgctgca
5521 cccgcgcttt atcgagcgtg tgcccgccgg cagccgcttt gactttcgcc tcaccgtgcg
5581 ccagcttgat ggcgacagcc ccgacctgct cgacaccctg ttggcgggcc tgaaaatgct
5641 ggagctggac gggctaggcg gcagcatttc ccgtgggtat ggcaaagtgc gctttgaagc
5701 gctgaccctc gacgggaaag atctccagcc gcgctttgag cagttacagc cgtttaaaca
5761 caccacgcag ggagccggat gaaagcttac ctggagatca aggagattac tctaacccca
5821 tcggccgtct taggggtttt ttgtcctgtg ttagctggag ggtataatac gactcactat
5881 agggagaccc gggataaagg aggtaaataa tgccctcccc cgttgccccc gccgccgcat
5941 ggcaatggct acgcctgcgg ctgctccccg acagcgcctt tgccaccgcg ctgcgcggtg
6001 acacgctgtt tggccagtta tgctggtatt tgcgcgaatc cctcggtgag gcgggcctga
6061 atgcgctgct ggcaggctac cacgaccagc gaccgttcgc ggtgatcagc gacccgatgc
6121 tggccgacca cctgccgcgc ccgcacctgc ccgaacaccg gctgggcttc gccgccgccg
6181 atgcccgcgc ccgcaaacaa cgtaaacagc aatgctggct gccgcttgcc ttcgctcacc
6241 agccgctggc cgagtggggc gcacacttaa ccgccgcccc tgagcaccac cagcaccgca
6301 gcgacgtgca gatgcacaac agcatcaacc gccagacact taccaccggt ggcgatgacg
6361 cgttcgcccc gtttggcagc gagcagcact ggtttgatgt ggacaccgcc tgggatctct
6721 tggcagacag cgccgccgtg ctggcccccg ttaccccgga taaccgcccg tttgtcggcc
6781 aggattagg cggcaatggc cgtctttcca ccgccctgcc acaaacggtg catcagggct
6841 atgcgccggt gatcccggtg cgctttcacc ataaggcaca gccacaatga tcatgatttc
6901 ttgtcgaact ggacagtagc agaaccgcta acgggggcga aggggttttt tgtgacatac
6961 gagctgattg aactaatacg actcactata gggagaggta ccataaagga ggtaaataat
7021 gacccgcgac cagacgccac gtcgccacac cgcctgtgac agcgacaaca tgcaatactt
7081 taccctcacc tgcctgtcgc cggtgcatgt cgccaccggc gacagcctga accccggtga
7141 atacctgata gacgaaaatg cactgtatga actggggcaa ggtggtctga gcccggcgct
7201 cacggcgaca cagcgcagcg agttgctcac tattctggag agcaatgacc ctgctctgcc
7261 gctgaccgtg cagcgttttc tggcacggga agcgggcaag ctgaaatacg ccgcccgccg
7321 gatgtgcccg ctgttacccg gcatcagccg ttattaccag tcacggctcg ggcaggtgat
7381 gcagaacgat accaaaaaca aaaagcagat gatcaaccag ctagagctga tgcgccacgt
7441 cggggccgca ctgggcgcgc cttatattcc cggatctacg ctcaaagggg ctatccgcac
7501 cgcactggtc agcgcgctca atcaaggcca gccactgcaa gccgagcgac gggaaaccga
7561 caaactgagc agcaaggtgg cgcaggatgc ggaacgccag ctactgggct ttgatacccg
7621 ccgcgacagc ccgcgatacc gcattgagca tgaccccttc cactggctac aggtggggga
7681 tgccgtaagc ccggcagagc acccgcccat gctggattac tggctggtac gccgccagcc
7741 gttcaaacgc accgagaagc aggacaacaa ggcggacaat atggaactgt cgccggttga
7801 atgccttaaa ccccggcaaa gcccgttgca ctgccagata accgtgaaaa cgccgcccac
7861 cgctctcgca ataaaaaacc cgcgcctgaa gcagtggctt ggcaaggtgt cgcaactggc
7921 gcaacaggtg aacaggataa ccctgccgca atgccatcac gagctggcat ggctggccga
7981 aaagcacatc ggcacggatg atgtgtacgc cccggccag aactgggtgg cgcagatgca
8041 gcaattactg cgccagttag acgacccgct acagcgcggc gaggcactct tgctgcgggt

| | |
|---|---|
| | 8101 gggcaaatac ggcggagcca tcagcaaaac cgtggcaggc tggcggcata tcgcccggct<br>8161 ggggagacaa ggcacgcgca ccacctacca cccggacgtc accacctgca cgctggcgct<br>8221 gccgcaagct gatgcgctga cacaggcgct gccctttggc tgggtactgt tgcaccagcc<br>8281 tgaccagccg gaggtgaccg agtttgtcgc cagccatcat gactggtgcc agcaacagca<br>8341 acagcggctt gacgcgcatc agcaacagca gcacacccac cgccagcaac gccagcagtt<br>8401 ggctcaggct cgtgaagagg aagcgcagcg gctggcagac aaagcccgcc agagcaaagc<br>8461 gcgccagtcc atcatgtcgc tggcagaaca actggcgagt gagcaaacgt ttcagcataa<br>8521 aaaccccaac ggcccgctgc gcggtcagtt agccacctgt gtgggttgcg ttgccactga<br>8581 aggctcagca gaagagaaag ccgagttgtg cacactgttt gacgacatcc tcaactattg<br>8641 gggcatcaaa cccggcaaag ataaaaagct cacggccttg aggaataagc tgttatgacc<br>8701 taggcgcttc aacggaacgg atcttacata tcgggggggt aggggttttt tgtctcggag<br>8761 accaagtagg gcataatacg actcactata gggagactat ggataaagga ggtaaataat<br>8821 gactaatcag gcaaaaaagt tatctagaat taatggtagg gagtttttaa aacagtcctt<br>8881 taatttacaa caacaactat tggcctctca attaaattta tcccgaacga ttacgcatga<br>8941 tggaacgatg ggggaggtta atgaaagtta ttttttgagt attatccgcc agtatttgcc<br>9001 tgaacgttac tcggttgacc ggggagttgt ggtggattca gaaggccaga ccagcgacca<br>9061 gatagatgca gtgatttttg accggcatta cacaccgaca ttattagacc aacaagggca<br>9121 caggtttatt ccggcagagg cggtgtacgc ggtactggag gttaaaccaa ccattaataa<br>9181 aacctacctt gaatatgcag ccgataaagc tgcatctgtc cgaaaattat atcgaaccag<br>9241 tacggtaata aaaaatattt acggtacggc caaaccggtc gaacatttcc cgatcgtagc<br>9301 aggtattgtg gcgattgatg ttgagtggca agacggactc ggaaaggcat ttactgaaaa<br>9361 tttgcaggct gtttccagcg atgaaaaccg aaaactggat tgcggtctgg cggtgtcggg<br>9421 cgcatgtttt gatagttatg atgaggaaat aaaaatcaga agcggtgaaa atgcattaat<br>9481 ctttttttctg ttccgtttgc tcggtaaatt gcaatcatta ggtacggtgc ccgcaattga<br>9541 ctggcgggtg tatatagata gtctggaata actcgagagg ttacagcctg cataatgtag<br>9601 cataacccct tggggcctct aaacgggtct tgaggggttt tttgtgccta tagtttgaag<br>9661 cagaatcgaa tttctgccat tcatccgctt attatcactt attcaggcgt agcaccaggc<br>9721 gtttaagggc accaataact gccttacaaa aaacccctag ccgcccgata agagcgggct<br>9781 aggggttcga gtaaaaaaaa ttacgccccg ccctgccact catcgcagta ctgttgtaat<br>9841 tcattaagca ttctgccgac atggaagcca tcacagacgg catgatgaac ctgaatcgcc<br>9901 agcggcatca gcaccttgtc gccttgcgta taatatttgc ccatcgtgaa aacgggggcg<br>9961 aagaagttgt ccatattggc cacgtttaaa tcaaaactgg tgaaactcac ccagggattg<br>10021 gctgaaacga aaaacatatt ctcaataaac cctttaggga aataggccag gttttcaccg<br>10081 taacacgcca catcttgcga atatatgtgt agaaactgcc ggaaatcgtc gtggtattca<br>10141 ctccagagcg atgaaaacgt ttcagtttgc tcatggaaaa cggtgtaaca agggtgaaca<br>10201 ctatcccata tcaccagctc accgtctttc attgccatac ggaactccgg gtgagcattc<br>10261 atcaggcggg caagaatgtg aataaaggcc ggataaaact tgtgcttatt tttctttacg<br>10321 gtctttaaaa aggccgtaat atccagctga acggtctggt tataggtaca ttgagcaact<br>10381 gactgaaatg cctcaaaatg ttctttacga tgccattggg atatatcaac ggtggtatat<br>10441 ccagtgattt ttttctccat tttagcttcc ttagctcctg aaaatctcga taactcaaaa<br>10501 aatacgcccg gtagtgatct tatttcatta tggtgaaagt tggaacctct tacgtgccga<br>10561 tcaaggtctc attttcgcca aaagttggcc cagggcttcc cggtatcaac agggacacca<br>10621 ggatttattt attctgcgaa gtgatcttcc gtcacaggta tttattcggc gcaaagtgcg<br>10681 tcgggtgatg ctgccaactt actgatttag tgtatgatgg tgtttttgag gtgctccagt<br>10741 ggcttctgtt tctatcagct gtccctcctg ttcagctact gacggggtgg tgcgtaacgg<br>10801 caaaagcacc gccggacatc agcgctagcg gagtgtatac tggcttacta tgttggcact<br>10861 gatgagggtg tcagtgaagt gcttcatgtg gcaggagaaa aaaggctgca ccggtgcgtc<br>10921 agcagaatat gtgatacagg atatattccg cttcctcgct cactgactcg ctacgctcgg<br>10981 tcgttcgact gcggcgagcg gaaatggctt acgaacgggg cggagatttc ctggaagatg<br>11041 ccaggaagat acttaacagg gaagtgagag ggccgcggca aagccgtttt tccataggct<br>11101 ccgcccccct gacaagcatc acgaaatctg acgctcaaat cagtggtggc gaaacccgac<br>11161 aggactataa agataccagg cgtttccccc tggcggctcc ctcgtgcgct ctcctgttcc<br>11221 tgcctttcgg tttaccggtg tcattccgct gttatggccg cgtttgtctc attccacgcc<br>11281 tgacactcag ttccgggtag gcagttcgct ccaagctgga ctgt |
| pTarget | 1 gtttgacagc ttatcatcga ctgcacggtg caccaatgct tctggcgtca ggcagccatc<br>61 ggaagctgtg gtatggctgt gcaggtcgta aatcactgca taattcgtgt cgctcaaggc<br>121 gcactcccgt tctggataat gttttttgcg ccgacatcat aacggttctg gcaaatattc<br>181 tgaaatgagc tgttgacaat taatcatccg gctcgtataa tgtgtggaat tgtgagcgga<br>241 taacaatttc acacaggaaa cagcgccgct gagaaaaagc gaagcggcac tgctctttaa<br>301 caatttatca gacaatctgt gtgggcactc gaccggaatt atcgattaac tttattatta<br>361 aaaattaaag aggtatatat taatgtatcg attaaataag gaggaataaa ccatgggggg<br>421 ttctcatcat catcatcatc atggtatggc tagcatgact ggtggacagc aaatgggtcg<br>481 ggatctgtac gacgatgacg ataaggatcc aacccttttc caagcttgca acttcattaa<br>541 gtggaacagg ttctgtaaat gcaagagcag catgcttcca aggcgaattc gaagcttggc<br>601 tgttttggcg gatgagagaa gattttcagc ctgatacaga ttaaatcaga acgcagaagc<br>661 ggtctgataa aacagaattt gcctggcggc agtagcgcgg tggtcccacc tgaccccatg<br>721 ccgaactcag aagtgaaacg ccgtagcgcc gatggtagtg tggggtctcc ccatgcgaga<br>781 gtagggaact gccaggcatc aaataaaacg aaaggctcag tcgaaagact gggcctttcg |

| | | |
|---|---|---|
| | 841 ttttatctgt tgtttgtcgg tgaacgctct cctgagtagg acaaatccgc cgggagcgga<br>901 tttgaacgtt gcgaagcaac ggcccggagg gtggcgggca ggacgcccgc cataaactgc<br>961 caggcatcaa attaagcaga aggccatcct gacggatggc ctttttgcgt ttctacaaac<br>1021 tcttttgttt atttttctaa atacattcaa atatgtatcc gctcatgaga caataaccct<br>1081 gataaatgct tcaataatat tgaaaaagga agagtatgag tattcaacat ttccgtgtcg<br>1141 cccttattcc cttttttgcg gcattttgcc ttcctgtttt tgctcaccca gaaacgctgg<br>1201 tgaaagtaaa agatgctgaa gatcagttgg gtgcacgagt gggttacatc gaactggatc<br>1261 tcaacagcgg taagatcctt gagagttttc gccccgaaga acgttttcca atgatgagca<br>1321 cttttaaagt tctgctatgt ggcgcggtat tatcccgtgt tgacgccggg caagagcaac<br>1381 tcggtcgccg catacactat tctcagaatg acttggttga gtactcacca gtcacagaaa<br>1441 agcatcttac ggatggcatg acagtaagag aattatgcag tgctgccata accatgagtg<br>1501 ataacactgc ggccaactta cttctgacaa cgatcggagg accgaaggag ctaaccgctt<br>1561 ttttgcacaa catgggggat catgtaactc gccttgatcg ttgggaaccg gagctgaatg<br>1621 aagccatacc aaacgacgag cgtgacacca cgatgcctgt agcaatggca acaacgttgc<br>1681 gcaaactatt aactggcgaa ctacttactc tagcttcccg gcaacaatta atagactgga<br>1741 tggaggcgga taaagttgca ggaccacttc tgcgctcggc ccttccggct ggctggttta<br>1801 ttgctgataa atctggagcc ggtgagcgtg ggtctcgcgg tatcattgca gcactggggc<br>1861 cagatggtaa gccctcccgt atcgtagtta tctacacgac ggggagtcag gcaactatgg<br>1921 atgaacgaaa tagacagatc gctgagatag gtgcctcact gattaagcat tggtaactgt<br>1981 cagaccaagt ttactcatat atactttaga ttgatttaaa acttcatttt taatttaaaa<br>2041 ggatctaggt gaagatcctt tttgataatc tcatgaccaa aatcccttaa cgtgagtttt<br>2101 cgttccactg agcgtcagac cccgtagaaa agatcaaagg atcttcttga gatccttttt<br>2161 ttctgcgcgt aatctgctgc ttgcaaacaa aaaaaccacc gctaccagcg gtggtttgtt<br>2221 tgccggatca agagctacca actctttttc cgaaggtaac tggcttcagc agagcgcaga<br>2281 taccaaatac tgtccttcta gtgtagccgt agttaggcca ccacttcaag aactctgtag<br>2341 caccgcctac atacctcgct ctgctaatcc tgttaccagt ggctgctgcc agtggcgata<br>2401 agtcgtgtct taccgggttg gactcaagac gatagttacc ggataaggcg cagcggtcgg<br>2461 gctgaacggg gggttcgtgc acacagccca gcttggagcg aacgacctac accgaactga<br>2521 gatacctaca gcgtgagcta tgagaaagcg ccacgcttcc cgaagggaga aaggcggaca<br>2581 ggtatccggt aagcggcagg gtcggaacag gagagcgcac gagggagctt ccagggggaa<br>2641 acgcctggta tctttatagt cctgtcgggt ttcgccacct ctgacttgag cgtcgatttt<br>2701 tgtgatgctc gtcagggggg cggagcctat ggaaaaacgc cagcaacgcg gcctttttac<br>2761 ggttcctggc cttttgctgg ccttttgctc acatgttctt tcctgcgtta tcccctgatt<br>2821 ctgtggataa ccgtattacc gcctttgagt gagctgatac cgctcgccgc agccgaacga<br>2881 ccgagcgcag cgagtcagtg agcgaggaag cggaagagcg cctgatgcgg tattttctcc<br>2941 ttacgcatct gtgcggtatt tcacaccgca tatggtgcac tctcagtaca atctgctctg<br>3001 atgccgcata gttaagccag tatacactcc gctatcgcta cgtgactggg tcatggctgc<br>3061 gccccgacac ccgccaacac ccgctgacgc gccctgacgg gcttgtctgc tcccggcatc<br>3121 cgcttacaga caagctgtga ccgtctccgg gagctgcatg tgtcagaggt tttcaccgtc<br>3181 atcaccgaaa cgcgcgaggc agcagatcaa ttcgcgcgcg aaggcgaagc ggcatgcatt<br>3241 tacgttgaca ccatcgaatg gtgcaaaacc tttcgcggta tggcatgata gcgcccggaa<br>3301 gagagtcaat tcagggtggt gaatgtgaaa ccagtaacgt tatacgatgt cgcagagtat<br>3361 gccggtgtct cttatcagac cgtttcccgc gtggtgaacc aggccagcca cgtttctgcg<br>3421 aaaacgcggg aaaaagtgga agcggcgatg gcggagctga attacattcc caaccgcgtg<br>3481 gcacaacaac tggcgggcaa acagtcgttg ctgattggcg ttgccacctc cagtctggcc<br>3541 ctgcacgcgc cgtcgcaaat tgtcgcggcg attaaatctc gcgccgatca actgggtgcc<br>3601 agcgtggtgg tgtcgatggt agaacgaagc ggcgtcgaag cctgtaaagc ggcggtgcac<br>3661 aatcttctcg cgcaacgcgt cagtgggctg atcattaact atccgctgga tgaccaggat<br>3721 gccattgctg tggaagctgc ctgcactaat gttccggcgt tatttcttga tgtctctgac<br>3781 cagacaccca tcaacagtat tattttctcc catgaagacg gtacgcgact gggcgtggag<br>3841 catctggtcg cattgggtca ccagcaaatc gcgctgttag cgggcccatt aagttctgtc<br>3901 tcggcgcgtc tgcgtctggc tggctggcat aaatatctca ctcgcaatca aattcagccg<br>3961 atagcggaac gggaaggcga ctggagtgcc atgtccggtt ttcaacaaac catgcaaatg<br>4021 ctgaatgagg gcatcgttcc cactgcgatg ctggttgcca acgatcagat ggcgctgggc<br>4081 gcaatgcgcg ccattaccga gtccgggctg cgcgttggtg cggatatctc ggtagtggga<br>4141 tacgacgata ccgaagacag ctcatgttat atcccgccgt taaccaccat caaacaggat<br>4201 tttcgcctgc tggggcaaac cagcgtggac cgcttgctgc aactctctca gggccaggcg<br>4261 gtgaagggca atcagctgtt gcccgtctca ctggtgaaaa gaaaaaccac cctggcgccc<br>4321 aatacgcaaa ccgcctctcc ccgcgcgttg gccgattcat taatgcagct ggcacgacag<br>4381 gtttcccgac tggaaagcgg gcagtgagcg caacgcaatt aatgtaagtt agcgcgaatt<br>4441 gatctg | |
| Name | Sequence (5’-3’) | Description |
| Gene fragments | | |

| | | |
|---|---|---|
| dPalm fragment | GCCAAAACCATCGGCCTGTCGCGCCAGATACACCTGTTTTTCA CTCTGTGGCTGCCGCATCTGTGCCGCAAAGACCCGCGTTTTGC CAACACCTACACCGTGTTCGCCGCAGCGGCCGCATTCTTTTTG ATTGGCCCGTGGCGCAGCCAGCAAACGCTGGCGCTAACAATGG CGCAGGACTTCGCGCGCTACAGCGGCCACAATCCGGCGCTGCA TTTTTCACTCGGGCTGGTGCAGGCCAAACCCGGCTACCCGGTG CGGGCGCTGGCGGCTCAGGCCGAAGCCGCTCTGAAGCAGGCCA AGCAACACCCCGGCAAGAACGCCATCTGCCTGTATAACGAGGT GATGGGCTGGCCGGAATATGACGCGTTACTGGCGTGCAGTGAG GAGCTGGCGCGCTGGCGCGAGCATGACGGCTACCCGCTCTCCT CCGGCCTGCTTTACCGGTTGCTGGCGCTCAGTGAACAGTCGGC GGAGGAGTCAGAAAAACCGCAGGCGGCACTGTGGCGCAGCCGG CTGGCCTATTTTTTACGTCGCAATCTGGTGGATAACGTGAAAG TCCCGACGAAAGAAAACCCGGCAGCGTTCCGCCATCAGCTTCA TTTGGAATTGTTTGAGAAACTTGAGCAGCACCTGAAACGGCAC CGCCAGCGTTACCGGGTGGCGCTTCAACGCCATCTCTACCACT ACCGCACGGTGCCGCGCGGCGAGTAACATAT | Mutates GGDD motif of Cas10 Palm2 domain to AAAA |
| dCsm3 fragment | ATGCAACTGAACAATATCCAGACGTTACGCGCCACTCTGGTG TGTGAAACCGGGTTACACATTGGCGGCGGCGACACCGCGTTG CAGATTGGCGGCATCGCCAGCGCCGTGGTGCGCCACCCGTTG ACCCAGCAACCTTACATTCCCGGCTCCAGCCTGAAAGGCAAA CTGCGCAGCCTGCTCGAATGGCGCGCTGGTGTGGTCGGCGAT ACCGAGGGCAAGGTGCTCAGTCATCAGGTTTACCAGCAACTG ATCGACGATAAAAAACAGGCACAGGCACTACAGGCACTGAAA ATCCTGCAACTGTTTGGCGTCAGCGGCGGCGACAAGCTTTCC GCCGAACAAGCCCAACAGATCGGCCCGACTCGCCTGTCCTTC TGGGATTGCGAATTTGACGAACACTGGCTGGCGCAGCAAGGC GGGCGCGTCCAGACCGAAGAGAAAGCGGAAAACTGTATTGAC CGCATCAGTGGCGTGGCGCTGCACCCGCGCTTTATCGAGCGT GTGCCCGCCGGCAGCCGCTTTGACTTTCGCCTCACCGTGCGC CAGCTTGATGGCGACAGCCCCGACCTGCTCGACACCCTGTTG GCGGGCCTGAAAATGCTGGAGCTGGACGGGCTAGGCGGCAGC ATTTCCCGTGGGTATGGCAAAGTGCGCTTTGAAGCGCTGACC CTCGACGGGAAAGATCTCCAGCCGCGCTTTGAGCAGTTACAG CCGTTTAAACACACCACGCAGGGAGCCGGATGA | Introduces D34A mutant to Csm3 |
| B01 fragment | TAATACGACTCACTATAGGGAGACCATGGGTCCTTACGGACG CTCCCTGACTGAAGGGATTAAGACTCCGGAGTCAGATGCACG ATGGTGTCTGTTTGAGGTCCTTACGGACGCTCCCTGACTGAAGG GATTAAGACCTCGAGGCTGTGGTCTAGACA | Cloning spacer B01, for expression of crRNA-B01 |
| S01 fragment | 5’TAATACGACTCACTATAGGGAGACCATGGGTCCTTACGGACGCT CCCTGACTGAAGGGATTAAGACACAGGAGTCAGATGCACGATGGT GTCTGTTTGAGGTCCTTACGGACGCTCCCTGACTGAAGGGATTAAG ACCTCGAGGCTGTGGTCTAGACA | Cloning spacer S01, for expression of crRNA-S01 |
| Oligonucleotides | | |

| | | |
|---|---|---|
| prCCP_077 | AATTCTGTTTTATCAGACCGTTATTCCAGACTATCTATATACACCCGCC | Rev primer to add pBAD homology to NucC |
| prCCP_077 | AATTCTGTTTTATCAGACCGTTATTCCAGACTATCTATATACACCCGCC | Rev primer to add pBAD homology to NucC |
| prCCP_078 | GGCTAACAGGAGGAATTAACATGACTAATCAGGCAAAAAAGTTATCTAGAAT | Fwd primer to add pBAD homology to NucC |
| prCCP_079 | GTTAATTCCTCCTGTTAGCCCAAAAAACGG | Rev primer to amplify region of interest from pBAD |
| prCCP_080 | CGGTCTGATAAAACAGAATTTGCCTGGC | Fwd primer to amplify region of interest from pBAD |
| prCCP_085 | GCAGCAGTGATTTTTGACCGGCATTACACA | Rev primer to introduce D83Q mutation in NucC |
| prCCP_086 | TATCTGGTCGCTGGTCTGGC | Fwd primer to introduce D83Q mutation in NucC |
| prCCP_087 | GCAGTTAAACCAACCATTAATAAAACCTACCTTGAATATG | Rev primer to introduce E114A mutation in NucC |
| prCCP_088 | CAGTACCGCGTACACCGCCTC | Fwd primer to introduce E114A mutation in NucC |
| prCCP_101 | CCACGCAGGGAGCCGGATGA | Rev primer to copy pACYC-Csmregion of interest for Gibson assembly |
| prCCP_102 | TGGATATTGTTCAGTTGCATTATTTACCTCCTTTATGA | Fwd primer to copy pACYC-Csm region of interest for Gibson assembly |
| prCCP_103 | GTGCCGCGCGGCGAGTAACATATG | Rev primer to copy pACYC-Csm region |

| | | |
|---|---|---|
| | | of interest for Gibson assembly |
| prCCP_104 | CGACAGGCCGATGGTTTTGGC | Fwd primer to copy pACYC-Csm region of interest for Gibson assembly |
| prCCP_109 | AGCGACGCAATAGATGCAGT | Fwd primer to introduce Q81A into pBAD NucC |
| prCCP_110 | GGTCTGGCCTTCTGAATCCA | Rev primer to introduce Q81A into pBAD NucC |
| prCCP_121 | CGTTCCGTTGAAGCGCCTAGGTC | Rev primer to make pACYC-Csm ΔNucC |
| prCCP_122 | GAATAACTCGAGAGGTTACAGCCTGCATAAT | Fwd primer to make pACYC-Csm ΔNucC |
| prCCP_185 | CACCGCGTTGGCGATTGGCGGCATCG | Fwd primer to make Q29A Csm3 |
| prCCP_186 | TCGCCGCCGCCAATGTGT | Rev primer to make Q29A Csm3 |
| prCCP_187 | CGACACCGCGGCGCAGATTGGCG | Fwd primer to make L28A Csm3 |
| prCCP_188 | CCGCCGCCAATGTGTAAC | Rev primer to make L28A Csm3 |
| prCCP_189 | GCGGGCGGCATCGACAGCG | Fwd primer to make I30A Csm3 |
| prCCP_190 | CTGCAACGCGGTGTCG | Rev primer to make I30A Csm3 |
| prCCP_191 | TGCGCTGTTAGCGAACCTGAAACCGC | Fwd primer to make H17A Cas10 |
| prCCP_192 | AAGGCAGCCACATGG | Rev primer to make H17A Cas10 |
| prIVT_001 | GAATTCTAATACGACTCACTATAGGGCAACTTCATTAAGTGGAA | Primer for overlap extension PCR to generate mismatch targets |

| | | |
|---|---|---|
| prIVT_002 | TACTTCCGCATTTACAGAACCTGTTCCACTTAATGAAGTTGC | Pairs with prIVT_001 to generate cognate target |
| prIVT_003 | TACTTCCGGATTTACAGAACCTGTTCCACTTAATGAAGTTGC | Pairs with prIVT_001 to generate +1 mismatch |
| prIVT_004 | TACTTCCGCTTTTACAGAACCTGTTCCACTTAATGAAGTTGC | Pairs with prIVT_001 to generate +2 mismatch |
| prIVT_005 | TACTTCCGCAATTACAGAACCTGTTCCACTTAATGAAGTTGC | Pairs with prIVT_001 to generate +3 mismatch |
| prIVT_006 | TACTTCCGCATATACAGAACCTGTTCCACTTAATGAAGTTGC | Pairs with prIVT_001 to generate +4 mismatch |
| prIVT_007 | TACTTCCGCATTAACAGAACCTGTTCCACTTAATGAAGTTGC | Pairs with prIVT_001 to generate +5 mismatch |
| prIVT_008 | TACTTCCGCATTTTCAGAACCTGTTCCACTTAATGAAGTTGC | Pairs with prIVT_001 to generate +6 mismatch |
| prIVT_009 | TACTTCCGCATTTAGAGAACCTGTTCCACTTAATGAAGTTGC | Pairs with prIVT_001 to generate +7 mismatch |
| prIVT_010 | TACTTCCGCATTTACTGAACCTGTTCCACTTAATGAAGTTGC | Pairs with prIVT_001 to generate +8 mismatch |
| prIVT_011 | TACTTCCGCATTTACACAACCTGTTCCACTTAATGAAGTTGC | Pairs with prIVT_001 to generate +9 mismatch |
| prIVT_012 | TACTTCCGCATTTACAGTACCTGTTCCACTTAATGAAGTTGC | Pairs with prIVT_001 to generate +10 mismatch |
| prIVT_013 | TACTTCCGCATTTACAGATCCTGTTCCACTTAATGAAGTTGC | Pairs with prIVT_001 to generate +11 mismatch |
| ssRNA-d02 | GCAACUUCAUUAAGUGGAACAGGUUCUGUAAAUGCGGAAGUA | |

| | | |
|---|---|---|
| Cognate | | |
| ssRNA-d03 Non-cognate | GCAACUUCAUUAAGUGGAACAGGUUCUGUAAAUGGUCUUAAU | |
| HBB target | CUCAAACAGACACCAUGGUGCAUCUGACUCCUGCGGAGAAGUCUGCCG | Synthetic RNA mimicking *HBB* transcript |
| A20T target | CUCAAACAGACACCAUGGUGCAUCUGACUCCUGAGGAGAAGUCUGCCG | Synthetic RNA mimicking the A20T transcript |
| Reporter DNA 1 | /56-FAM/TCAAAGGCGCCCTTGGTACATACACGATTT | Cleaved by NucC in fluorogenic assay |
| Reporter DNA 2 | AAATCGTGTATGTACCAAGGGCGCCTTTGA/31ABkFQ/ | Cleaved by NucC in fluorogenic assay |

**Table S2. Buffers used in Cas10-Csm purification**

| Name | Reagent |
|---|---|
| **Phosphate lysis buffer** | 100 mM $NaH_2PO_4$ pH 8.0 |
| | 600 mM NaCl |
| | 20 mM imidazole |
| | 1 mg/mL lysozyme |
| | 2 mM PMSF |
| | 0.1% (v/v) triton X-100 |
| | 1% v/v glycerol |
| | 1 mM DTT |
| | 500 µM EDTA |
| **Phosphate wash buffer 1** | 100 mM $NaH_2PO_4$ |
| | 150 mM NaCl |
| | 1 mM DTT |
| | 1% (v/v) glycerol |
| | 20 mM imidazole |
| **Phosphate wash buffer 2** | Phosphate wash buffer 1 with 40 mM imidazole |
| **Phosphate elution buffer 1** | Phosphate wash buffer 1 with 100 mM imidazole |
| **Phosphate elution buffer 2** | Phosphate wash buffer 1 with 250 mM imidazole |
| **Tris lysis buffer** | 50 mM Tris-HCl pH 8.0 |
| | 300 mM NaCl |
| | 20 mM imidazole |
| | 1 mg/ml lysozyme |
| | 1 mM PMSF |
| | 0.1% (v/v) triton X-100 |
| **Tris wash buffer** | Tris lysis buffer with 40 mM imidazole and 10% (v/v) glycerol |
| **Tris elution buffer 1** | Tris lysis buffer with 100 mM imidazole and 10% (v/v) glycerol |
| **Tris elution buffer 2** | Tris lysis buffer with 250 mM imidazole and 10% (v/v) glycerol |
| **Storage buffer** | 50 mM Tris HCl pH 8.0 |
| | 20 mM NaCl |
| | 5% (v/v) glycerol |
| **Reaction buffer 1** | 50 mM Tris HCl pH 8.0 |
| | 150 mM $NH_4Cl$ |
| | 5% (v/v) glycerol |
| | 10 mM $MgCl_2$ |
| **Reaction buffer 2** | 50 mM Tris HCl pH 8.0 |
| | 150 mM NaCl |
| | 5% (v/v) glycerol |
| | 10 mM $MgCl_2$ |
| **Reaction buffer 3** | 12.5 mM Tris HCl pH 8.0 |
| | 20 mM NaCl |
| | 10 mM $MgCl_2$ |
| | 10% v/v glycerol |
| **EM buffer** | 50 mM Tris HCl pH 8.0 |
| | 100 mM NaCl |
| | 1 mM $MgSO_4$ |
| | 4 mM $CaCl_1$ |

**Table S3. Mass spectrometry identification of proteins in SmCas10-Csm ultracentrifugation fractions.**

| Protein | Theoretical mass (kDa) | UNIPARC Identifiers | Unique peptide counts | PSMs[a] | Sum PEP score |
|---|---|---|---|---|---|
| **Cas10** | 90.2 | UPI0003926B6B | 29 | 358 | 80.165 |
| **Csm5** | 63 | UPI0003921A6A | 21 | 431 | 80.144 |
| **ArnA** | 74.2 | UPI0000000F69 | 26 | 96 | 64.091 |
| **Csm3** | 27.2 | UPI00039276A8 | 19 | 892 | 53.839 |
| **Csm2** | 15.8 | UPI000392639F | 13 | 583 | 39.57 |
| **Csm4** | 35.7 | UPI001F2BC94D | 10 | 219 | 39.294 |
| **GroEL** | 57.3 | UPI0000000ED4 | 14 | 32 | 30.367 |
| **AdhE** | 96.1 | UPI0000000054 | 16 | 37 | 29.725 |
| **SlyD** | 20.8 | UPI0000135A43 | 5 | 19 | 25.797 |

[a] All proteins comprising 1% or more of total protein as determined by number PSM are reported.
PSM - Peptide spectrum match
PEP - Posterior error probability for peptide spectrum matches, PEP score is the negative logarithm of the PEP value

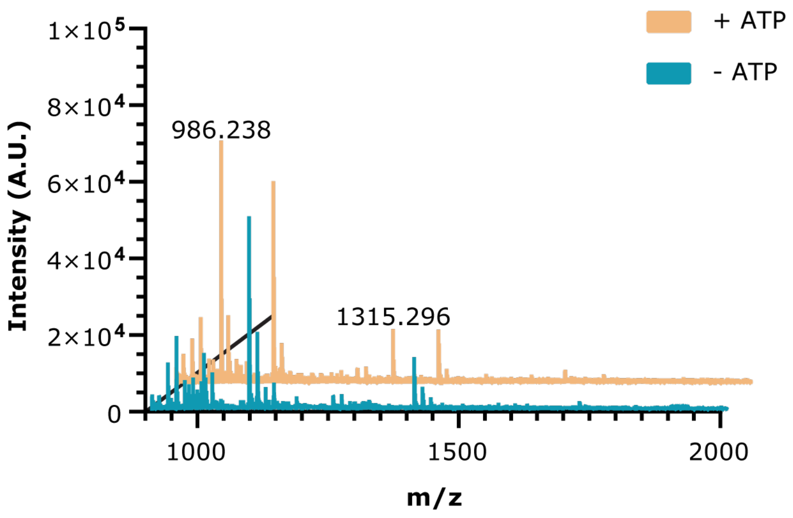


**Figure S3. MALDI mass spectrometry of cOA synthesis reactions plus and minus ATP.**

**Table S4. Cryo-EM Statistics for Data Collection and Model Quality**

| **Data collection and processing** | **Cas10-Csm unbound to target RNA**<br>**9ZS2**<br>**EMD-74675** | **Cas10-Csm bound to target RNA**<br><br>**9ZS4**<br>**EMD-74688** |
|---|---|---|
| Magnification | 150,000 | 150,000 |
| Voltage (kV) | 200 | 200 |
| Electron exposure ($e^-$/ Å$^2$) | 40.0 | 40.0 |
| Defocus range (μm) | 0.1-3.5 | 0.1-3.0 |
| Pixel size (Å) | 0.94 | 0.94 |
| Initial particles (no.) | 593,739 | 324,865 |
| Final particles (no.) | 42,471 | 129,762 |
| Map resolution (Å) | 4.4 | 3.8 |
| FSC threshold | 0.143 | 0.143 |
| Map sharpening B factor (Å$^2$) | 118.6 | 110.9 |
| **Refinement** | | |
| Model resolution (Å) | 4.4 | 3.8 |
| FSC threshold | 0.143 | 0.143 |
| Model composition | | |
| Nonhydrogen atoms | 18,543 | 23,918 |
| Protein residues | 2308 | 2867 |
| RNA residues | 30 | 64 |
| Bonds (RMSD) | | |
| Bond lengths (Å) | 0.003 | 0.003 |
| Bond angles (°) | 0.770 | 0.742 |
| Validation | | |
| Molprobity score | 1.99 | 2.00 |
| Clashscore | 15.68 | 9.85 |
| Ramachandran plot (%) | | |
| Outliers | 0.22 | 0.00 |
| Allowed | 4.02 | 3.34 |
| Favored | 95.76 | 96.66 |
| B factors, mean (Å$^2$) | | |
| Protein | 420.62 | 203.32 |
| RNA | 313.83 | 148.32 |

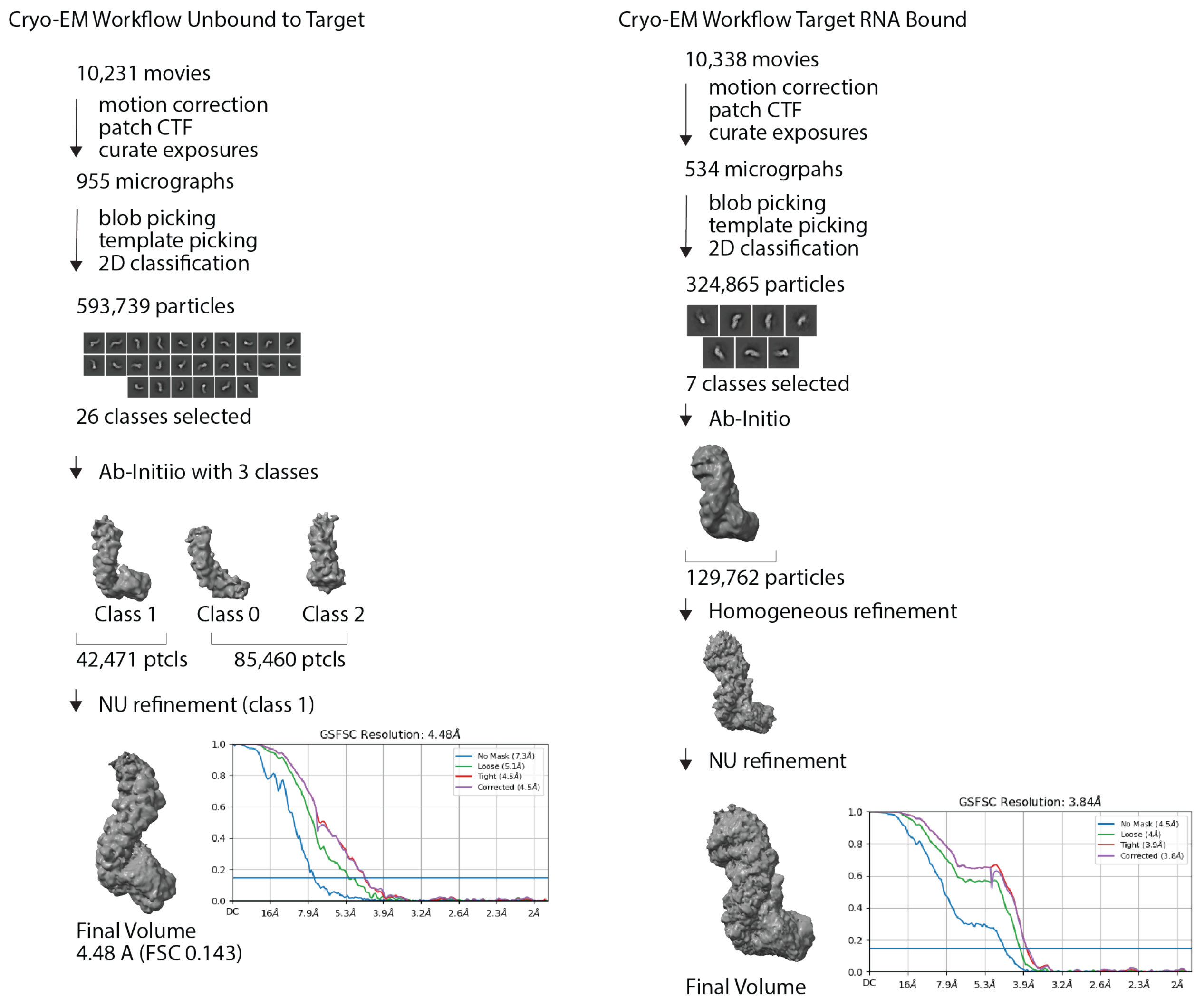


**Figure S4. Workflow for single particle reconstructions of SmCas10-Csm by cryo-EM.**

### Table S5. Fluorescence signal for titrations of wild type *HBB* RNA and variant

crRNA-B01/wt *HBB* RNA

| Concentration | Raw fluorescence | | Background subtracted intensity | I/σ(I) |
|---|---|---|---|---|
| | **Mean** | **σ** | **Mean** | |
| **1 uM** | 11459 | 541 | 8926 | 16.5 |
| **100 nM** | 10698 | 783 | 8166 | 10.4 |
| **10 nM** | 11084 | 1163 | 8551 | 7.4 |
| **1 nM** | 4036 | 267 | 1504 | 5.6 |
| **100 pM** | 2624 | 203 | 92 | 0.5 |
| **10 pm** | 2520 | 147 | -12 | -0.1 |
| **1 pm** | 2382 | 215 | -151 | -0.7 |
| | Blank mean 2533 | Blank s = 197 | | |

crRNA-B01/A20T RNA

| Concentration | Raw fluorescence | | Background subtracted intensity | I/σ(I) |
|---|---|---|---|---|
| | **Mean** | **σ** | **Mean** | |
| **1 uM** | 10496 | 769 | 7986 | 10.4 |
| **100 nM** | 4219 | 492 | 1709 | 3.5 |
| **10 nM** | 2708 | 152 | 198 | 1.3 |
| **1 nM** | 2397 | 168 | -113 | -0.7 |
| **100 pM** | 2624 | 120 | 115 | 1.0 |
| **10 pm** | 2066 | 105 | -444 | -4.2 |
| **1 pm** | 2608 | 122 | 98 | 0.8 |
| | Blank mean 2509 | Blank s = 266 | | |

crRNA-S01/A20T RNA

| Concentration | Raw Fluorescence | | Background subtracted intensity | I/σ(I) |
|---|---|---|---|---|
| | **Mean** | **σ** | **Mean** | |
| **1 uM** | 15129 | 162 | 14013 | 86.6 |
| **100 nM** | 15069 | 345 | 13952 | 40.5 |
| **10 nM** | 13574 | 2133 | 12458 | 5.8 |
| **1 nM** | 9623 | 1836 | 8507 | 4.6 |
| **100 pM** | 1794 | 390 | 678 | 1.7 |
| **10 pm** | 1217 | 151 | 101 | 0.7 |
| **1 pm** | 1198 | 187 | 82 | 0.4 |
| | Blank mean 1116 | Blank s = 191 | | |

crRNA-S01/wt *HBB* RNA

| Concentration | Raw Fluorescence | | Background subtracted intensity | I/σ(I) |
|---|---|---|---|---|
| **1** | **Mean** | **σ** | **Mean** | |
| **1 uM** | 5959 | 492 | 4760 | 9.7 |
| **100 nM** | 6402 | 711 | 5204 | 7.3 |
| **10 nM** | 1725 | 233 | 527 | 2.3 |
| **1 nM** | 1197 | 169 | -1 | 0.0 |
| **100 pM** | 1074 | 253 | -124 | -0.5 |
| **10 pm** | 1216 | 134 | 18 | 0.1 |
| **1 pm** | 1406 | 460 | 208 | 0.5 |
| | Blank mean 1198 | Blank s = 76 | | |

*I/σ(I), mean of background subtracted intensity divided by standard deviation of this mean.*

**Table S6. Fluorescence signal for SNP detection in contrived samples.**

**crRNA-B01**

| **Targets** | **Raw fluorescence** | | **Background subtracted intensity** | **I/ σ(I)** |
|---|---|---|---|---|
| | **Mean** | **σ** | **Mean** | |
| **HBB** | 13314 | 293 | 11585 | 39.5 |
| **HBB / A20T** | 11645 | 131 | 9916 | 75.6 |
| **A20T** | 4292 | 566 | 2564 | 4.5 |
| | Blank mean 1729 | Blank s = 143 | | |

**crRNA-S01**

| **Targets** | **Raw fluorescence** | | **Background subtracted intensity** | **I/ σ(I)** |
|---|---|---|---|---|
| | **Mean** | **σ** | **Mean** | |
| **HBB** | 3702 | 451 | 174 | 0.4 |
| **HBB / A20T** | 12817 | 360 | 9288 | 25.8 |
| **A20T** | 12010 | 88 | 8481 | 96.7 |
| | Blank mean 3529 | Blank s = 414 | | |
